# Deep Learning-Inferred Multiplex ImmunoFluorescence for IHC Image Quantification

**DOI:** 10.1101/2021.05.01.442219

**Authors:** Parmida Ghahremani, Yanyun Li, Arie Kaufman, Rami Vanguri, Noah Greenwald, Michael Angelo, Travis J. Hollmann, Saad Nadeem

**Affiliations:** Department of Computer Science, Stony Brook University, Stony Brook, NY, USA; Department of Pathology, Stanford University, Stanford, CA, USA; Department of Pathology, Memorial Sloan Kettering Cancer Center, New York, NY, USA; Department of Medical Physics, Memorial Sloan Kettering Cancer Center, New York, NY, USA

**Keywords:** Multitask Learning, Multiplex ImmunoFluoresence, Immunohistochemisty

## Abstract

Reporting biomarkers assessed by routine immunohistochemical (IHC) staining of tissue is broadly used in diagnostic pathology laboratories for patient care. To date, clinical reporting is predominantly qualitative or semi-quantitative. By creating a multitask deep learning framework referred to as DeepLIIF, we present a single-step solution to stain deconvolution/separation, cell segmentation, and quantitative single-cell IHC scoring. Leveraging a unique *de novo* dataset of co-registered IHC and multiplex immunofluorescence (mpIF) staining of the same slides, we segment and translate low-cost and prevalent IHC slides to more expensive-yet-informative mpIF images, while simultaneously providing the essential ground truth for the superimposed brightfield IHC channels. Moreover, a new nuclear-envelop stain, LAP2beta, with high (>95%) cell coverage is introduced to improve cell delineation/segmentation and protein expression quantification on IHC slides. By simultaneously translating input IHC images to clean/separated mpIF channels and performing cell segmentation/classification, we show that our model trained on clean IHC Ki67 data can generalize to more noisy and artifact-ridden images as well as other nuclear and non-nuclear markers such as CD3, CD8, BCL2, BCL6, MYC, MUM1, CD10, and TP53. We thoroughly evaluate our method on publicly available benchmark datasets as well as against pathologists’ semi-quantitative scoring. The code, the pre-trained models, along with easy-to-run containerized docker files as well as Google CoLab project are available at https://github.com/nadeemlab/deepliif.

## Introduction

The assessment of protein expression using immunohistochemical staining of tissue sections on glass slides is critical for guiding clinical decision-making in several diagnostic clinical scenarios, including cancer classification, residual disease detection, and even mutation detection (BRAFV600E and NRASQ61R). Standard brightfield chromogenic IHC staining, while high throughput, has a narrow dynamic range and results in superimposed channels with high chromogen/stain overlap, requiring specialized digital stain deconvolution/separation, e.g. (1), as an essential preprocessing step in both state-of-the-art research as well as commercial IHC quantification algorithms. Stain deconvolution is an open problem requiring extensive hyper-parameter tuning (on per-case basis) or (highly-error prone and time consuming) manual labeling of different cell types (2, 3), but still results in sub-optimal color separation in regions of high chromogen overlap.

As opposed to standard brightfield IHC staining, multiplex immunofluorescence (mpIF) staining provides the opportunity to examine panels of several markers individually (without requiring stain deconvolution) or simultaneously as a composite permitting accurate co-localization, stain standardization, more objective scoring, and cut-offs for all the markers’ values (especially in low-expression regions, which are difficult to assess on IHC stained slides and can be misconstrued as negative due to weak staining that can be masked by the hematoxylin counterstain) (4, 5). Moreover, in a recent meta-analysis (6), mpIF was shown to have a higher diagnostic prediction accuracy (at par with multimodal crossplatform composite approaches) than IHC scoring, tumor mutational burden, or gene expression profiling. However, mpIF assays are expensive and not widely available. This can lead to a unique opportunity to leverage the advantages of mpIF to improve the explainability and interpretability of the conventional IHCs using recent deep learning breakthroughs. Current deep learning methods for scoring IHCs rely solely on the error-prone manual annotations (unclear cell boundaries, overlapping cells, and challenging assessment of low-expression regions) rather *than on co-registered highdimensional imaging of the same tissue samples (that can provide essential ground truth for the superimposed brightfield IHC channels).* Therefore, we present a new multitask deep learning algorithm that leverages a unique coregistered IHC and mpIF training data of the same slides to simultaneously translate low-cost/prevalent IHC images to high-cost and more informative mpIF representations (creating a Deep-Learning-Inferred IF image), accurately autosegment relevant cells, and quantify protein expression for more accurate and reproducible IHC quantification; using multitask learning (7) to train models to perform a variety of tasks rather than one narrowly defined task makes them more generally useful and robust. Specifically, once trained, DeepLIIF takes only IHC image as input (e.g., Ki67 protein IHC as a brown Ki67 stain with standard hematoxylin nuclear counterstain) and completely bypassing stain deconvolution, produces/generates corresponding hematoxylin, mpIF nuclear (DAPI), mpIF protein (e.g., Ki67), mpIF LAP2Beta (a new nuclear envelop stain with *>* 95% cell coverage to better separate touching/overlapping cells) channels and segmented/classified cells (e.g., Ki67+ and Ki67-cell masks for estimating Ki67 proliferation index which is an important clinical prognostic metric across several cancer types), as shown in Figure 1. Moreover, DeepLIIF trained just on clean IHC Ki67 images generalizes to more noisy and artifact-ridden images as well as other nuclear and non-nuclear markers such as CD3, CD8, BCL2, BCL6, MYC, MUM1, CD10, and TP53. Example IHC images stained with different markers along with the DeepLIIF inferred modalities and segmented/classified nuclear masks are also shown in Figure 1. *In essence, DeepLIIF presents a single-step solution to stain deconvoluion, cell segmentation, and quantitative single-cell IHC scoring. Additionally, our co-registered mpIF data, for the first time, creates an orthogonal dataset to confirm and further specify the target brightfield IHC staining characteristics.*

**Figure 1.**
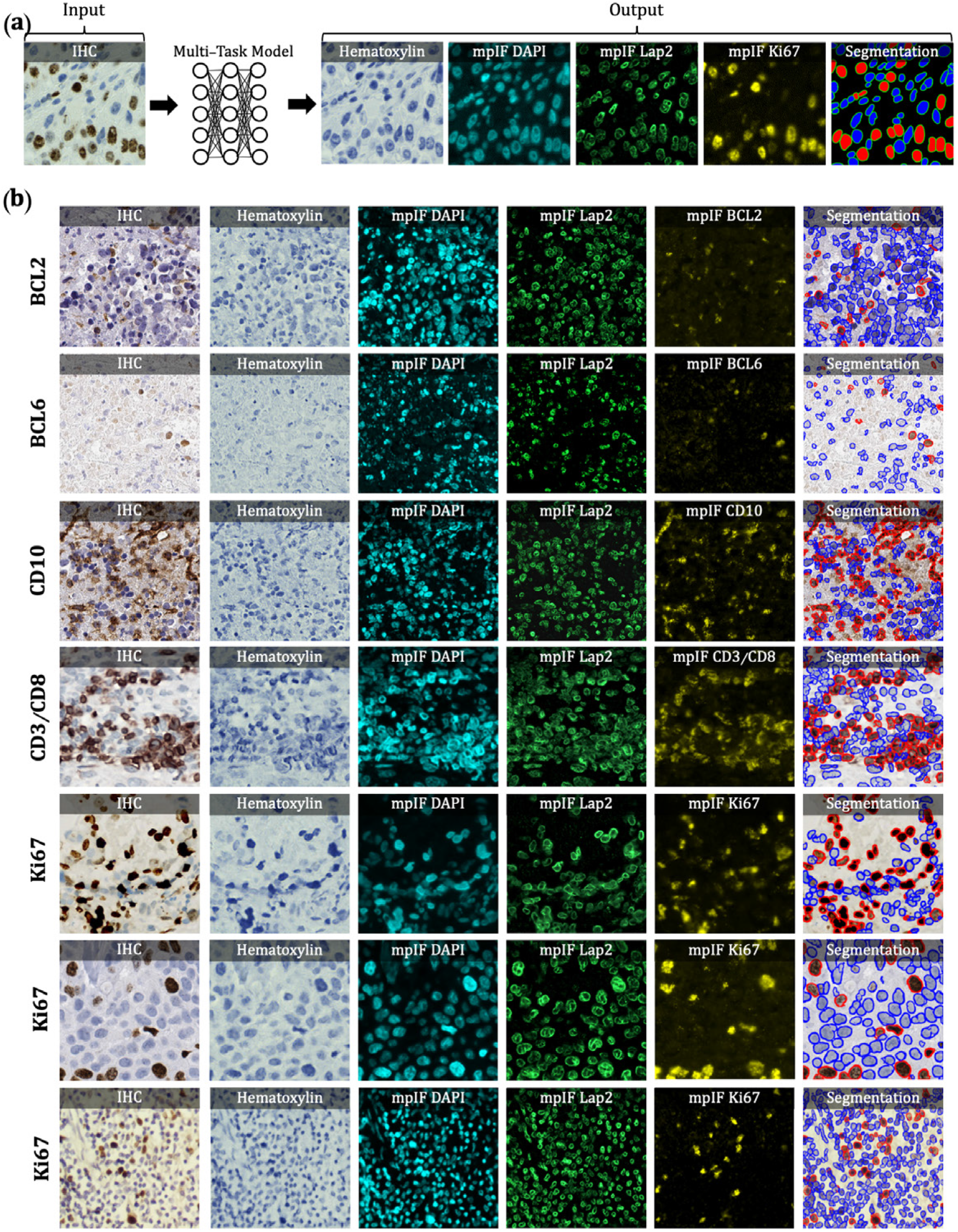
Overview of DeepLIIF pipeline and sample input IHCs (different brown/DAB markers – BCL2, BCL6, CD10, CD3/CD8, Ki67) with corresponding DeepLIIF-generated hematoxylin/mpIF modalities and classified (positive (red) and negative (blue) cell) segmentation masks. (a) Overview of DeepLIIF. Given an IHC input, our multitask deep learning framework simultaneously infers corresponding Hematoxylin channel, mpIF DAPI, mpIF protein expression (Ki67, CD3, CD8, etc.), and the positive/negative protein cell segmentation, baking explainability and interpretability into the model itself rather than relying on coarse activation/attention maps. In the segmentation mask, the red cells denote cells with positive protein expression (brown/DAB cells in the input IHC), whereas blue cells represent negative cells (blue cells in the input IHC). (b) Example DeepLIIF-generated hematoxylin/mpIF modalities and segmentation masks for different IHC markers. DeepLIIF, trained on clean IHC Ki67 nuclear marker images, can generalize to noisier as well as other IHC nuclear/cytoplasmic marker images.

## Results

In this section, we evaluate the performance of DeepLIIF on cell segmentation and classification tasks. We evaluated the performance of our model and other state-of-the-art methods using pixel accuracy (PixAcc) computed from the number of true positives, TP, false positives, FP and false negatives, FN, as 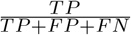, Dice Score as 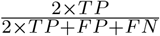, and IOU as the class-wise intersection over the union. We compute these metrics for each class, including negative and positive, and compute the average value of both classes for each metric. A pixel is counted as TP if it is segmented and classified correctly. A pixel is considered FP if it is falsely segmented as the foreground of the corresponding class. A pixel is counted as FN if it is falsely detected as the background of the corresponding class. For example, assuming the model segments a pixel as a pixel of a negative cell (blue), but in the groundtruth mask, it is marked as positive (red). Since there is no corresponding pixel in the foreground of the ground-truth mask of the negative class, it is considered FP for the negative class and FN for the positive class, as there is no marked corresponding pixel in the foreground of the predicted mask of the positive class. We also evaluate our model against other methods using Aggregated Jaccard Index (AJI) which is an object-level metric (8), defined as 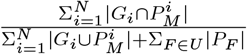. Considering that the goal is an accurate interpretation of IHC staining results, we compute the difference between the IHC quantification percentage of the predicted mask and the real mask, as shown in Figure 2.

**Figure 2.**
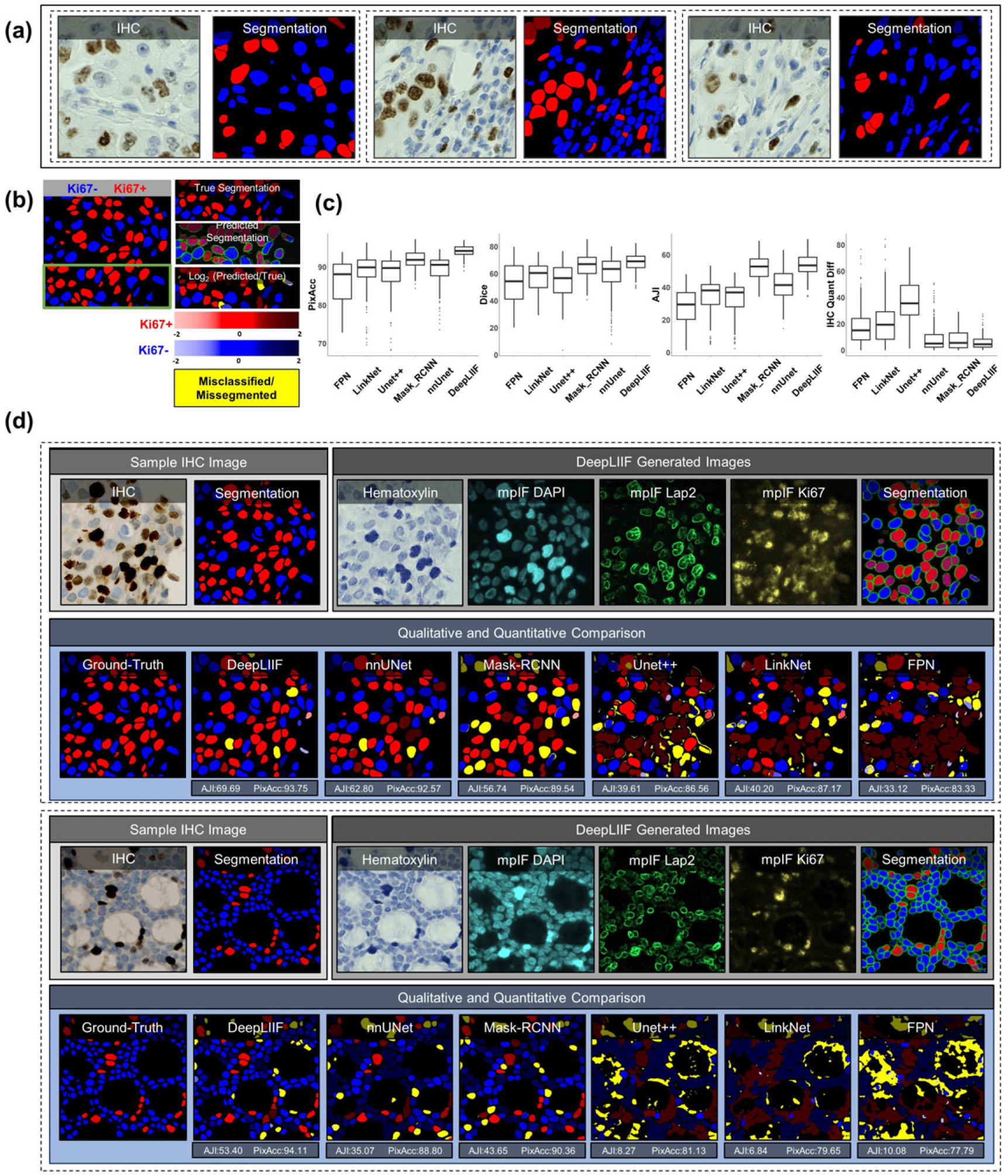
Qualitative and quantitative analysis of DeepLIIF against state-of-the-art semantic segmentation models tested on BC Dataset (9). (a) Three example images from our training set. (b) A segmentation mask showing Ki67- and Ki67+ cell representation, along with a visual segmentation and classification accuracy. Predicted classes are shown in different colors where blue represents Ki67- and red represents Ki67+ cells, and the hue is set using the *log2* of the ratio between the predicted area and ground-truth area. Cells with too large areas are shown in dark colors, and cells with too small areas are shown in a light color. For example, if the model correctly classifies a cell as Ki67+, but the predicted cell area is too large, the cell is colored in dark red. If there is no cell in the ground-truth mask corresponding to a predicted cell, the predicted cell is shown in yellow, which means that the cell is misclassified (cell segmented correctly but classified wrongly) or missegmented (no cell in the segmented cell area). (c) The accuracy of the segmentation and classification is measured by getting the average of Dice score, Pixel Accuracy, absolute value of IHC Quantification difference between the predicted segmentation mask of each class and the ground-truth mask of the corresponding class (0 indicates no agreement and 100 indicates perfect agreement). Evaluation of all scores shows that DeepLIIF outperforms all state-of-the-art models. (d) As mentioned earlier, DeepLIIF generalizes across different tissue types and imaging platforms. Two example images from the BC Dataset (9) along with the inferred modalities and generated classified segmentation masks are shown in the top rows where the ground-truth mask and segmentation masks of five state-of-the-art models are shown in the second row. The mean IOU and Pixel Accuracy are given for each model in the box below the image.

To compare our model with state-of-the-art models, we use three different datasets. 1) We evaluate all models on our internal test set, including 600 images of size 512 × 512 and 40x magnification from bladder carcinoma and non-small cell lung carcinoma slides. 2) We randomly selected and segmented 41 images of size 640 × 640 from recently released BCDataset (9) which contains Ki67 stained sections of breast carcinoma from scanned whole slide images with manual Ki67+ and Ki67-cell centroid annotations (targeting cell detection as opposed to cell instance segmentation task), created from consensus of 10 pathologists. We split these tiles into 164 images of size 512 × 512; the test set varies widely in the density of tumor cells and the Ki67 index. 3) We also tested our model and others on a publicly available CD3 and CD8 IHC NuClick Dataset (10). We used the training set of BC Dataset containing 671 IHC patches of size 256 × 256, extracted from LYON19 dataset (11). LYON19 (11) is a Grand Challenge to provide a dataset and an evolution platform to benchmark existing algorithms for lymphocyte detection in IHC stained specimens. The dataset contains IHC images of breast, colon, and prostate stained with an antibody against CD3 or CD8.

Trained on clean lung and bladder images stained with Ki67 marker, DeepLIIF generalizes well to other markers. We trained state-of-the-art segmentation networks, including *FPN* (12), *LinkNet* (13), *Mask_RCNN* (14), *Unet++* (15), and *nnU-Net* (16) on our training set (described in Section Training Data) using the IHC images as the input and generating the colored segmentation mask representing normal cells and lymphocytes. DeepLIIF outperformed previous models trained and tested on the same data on all three metrics. We trained and tested all models on a desktop with an NVIDIA Quadro RTX 6000 GPU, which was also used for all implementations.

We compare the DeepLIIF model’s performance against state-of-the-art models on the test set obtained from BC-Dataset (9). The results were analyzed both qualitatively and quantitatively, as shown in Figure 2. All models are trained and validated on the same training set as the DeepLIIF model. Application of DeepLIIF to the BC Dataset resulted in a pixel accuracy of 94.18%, Dice score of 68.15%, IOU of 53.20%, AJI of 53.48%, and IHC quantification difference of 6.07%, and outperformed *Mask_RCNN* with pixel accuracy of 91.95%, IOU of 66.16%, Dice Score of 51.16%, AJI of 52.36%, and IHC quantification difference of 8.42%, *nnUnet* with pixel accuracy of 89.24%, Dice Score of 58.69%, IOU of 43.44%, AJI of 41.31%, and IHC quantification difference of 9.84%, *UNet++* with pixel accuracy of 87.99%, Dice Score of 54.91%, IOU of 39.47%, AJI of 32.53%, and IHC quantification difference of 36.67%, *LinkNet* with pixel accuracy of 88.59%, Dice score of 33.64%, IOU of 41.63%, AJI of 33.64%, and IHC quantification difference of 21.57%, and *FPN* with pixel accuracy of 85.78%, Dice score of 52.92%, IOU of 38.04%, AJI of 27.71%, and IHC quantification difference of 17.94%, while maintaining lower standard deviation on all metrics. We also performed a significance test to show that DeepLIIF significantly outperforms other models. As mentioned earlier, all models are trained and tested on the exact same dataset, meaning that the data is paired. Therefore, we perform a paired Wilcoxon rank-sum test, where a p-value of 5% or lower is considered statistically significant. All tests are two-sided, and the assumption of normally distributed data was tested using a Shapiro-Wilk test. The computed p-values of all metrics show that DeepLIIF significantly outperforms the state-of-the-art models.

We used pixel-level accuracy metrics for the primary evaluation, as we are formulating the IHC quantification problem as cell instance segmentation/classification. However, since DeepLIIF is capable of separating the touching nuclei, we also performed a cell-level analysis of DeepLIIF against cell centroid detection approaches. *U-CSRNet* (9), for example, detects and classifies cells without performing cell instance segmentation. Most of these approaches use crowd-counting techniques to find cell centroids. The major hurdle in evaluating these techniques is the variance in detected cell centroids. We trained *FCRN_A* (17), *FCRN_B* (17), *Deeplab_Xeption* (18), *SC_CNN* (19), *CSRNet* (20), *U-CSRNet* (9) using our training set (the centroids of our individual cell segmentation masks are used as detection masks). Most of these approaches failed in detecting and classifying cells on the BCData testing set, and the rest detected centroids far from the ground-truth centroids. As a result, we resorted to comparing the performance of DeepLIIF (trained on our training set) with these models trained on the training set of the BCDataset and tested on the testing set of the BCData. As shown in Exteneded Data Figure 1, even though our model was trained on a completely different dataset from the testing set, it has better performance than the detection models that were trained on the same training set of the test dataset. The results show that, unlike DeepLIIF, the detection models are not robust across different datasets, staining techniques, and tissue/cancer types.

As was mentioned earlier, our model generalizes well to segment/classify cells stained with different markers, including CD3/CD8. We compare the performance of our trained model against other trained models on the training set of the NuClick dataset (21). The comparative analysis is shown in Figure 3. The DeepLIIF model outperformed other models on segmenting and classifying CD3/CD8+ cells (tumorinfiltrating lymphocytes or TILs) on all three metrics.

**Figure 3.**
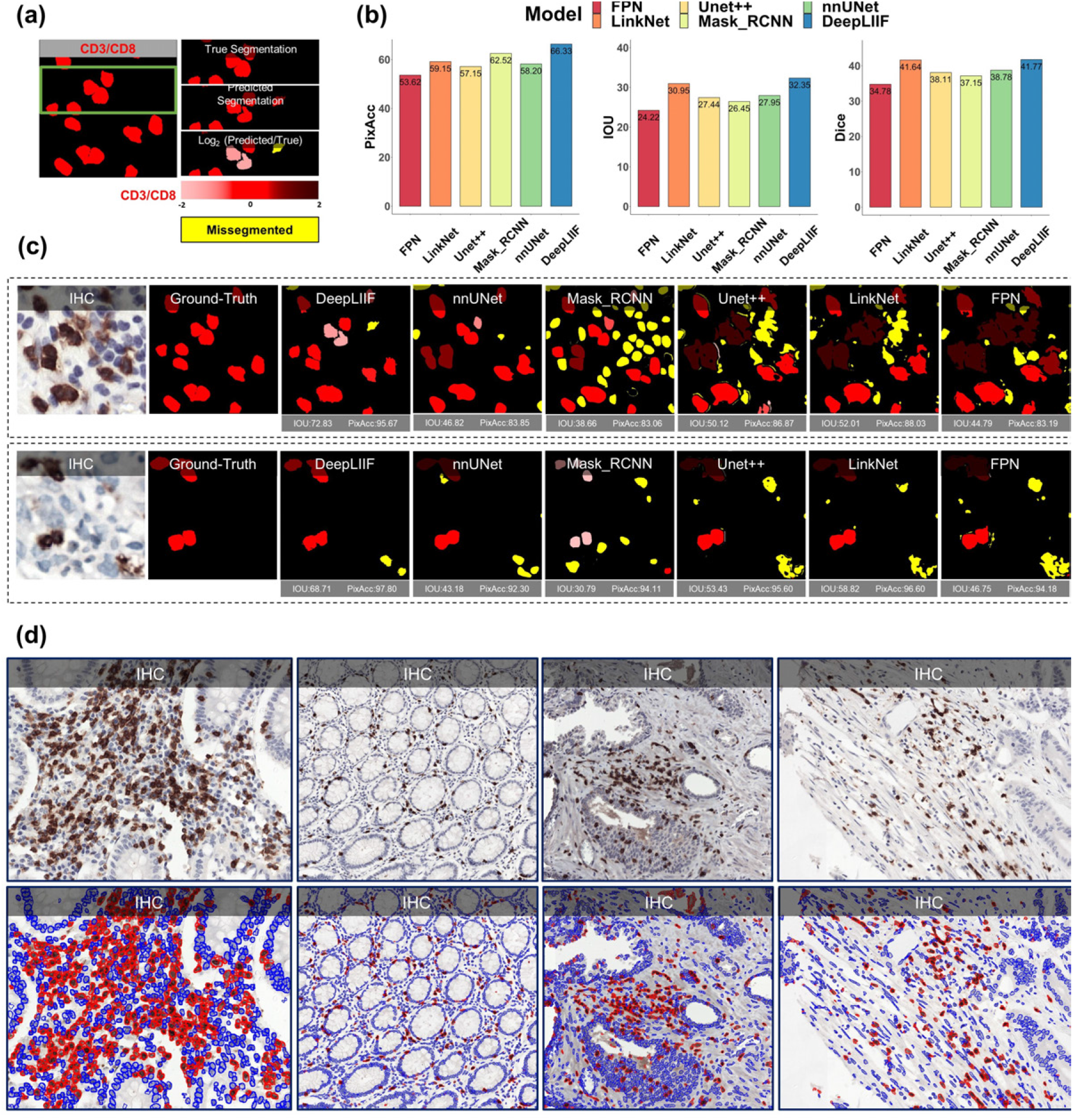
Qualitative and quantitative analysis of DeepLIIF against state-of-the-art semantic segmentation models tested on NuClick Dataset (21) and four sample images from the LYON19 challenge dataset (11). (a) A segmentation mask showing CD3/CD8+ cells, along with a visual segmentation and classification accuracy. Predicted CD3/CD8+ cells are shown in red color, and the hue is set using the *log*_2_ of the ratio between the predicted area and ground-truth area. Cells with too large areas are shown in dark colors, and cells with too small areas are shown in a light color. For example, if the model correctly classifies a cell as CD3/CD8+, but the predicted cell area is too large, the cell is colored in dark red. If there is no cell in the ground-truth mask corresponding to a predicted cell, the predicted cell is shown in yellow, which means that the cell is missegmented (no corresponding ground-truth cell in the segmented cell area). (b) The accuracy of the segmentation and classification is measured by getting the average of Dice score, Pixel Accuracy, and IOU (intersection over union) between the predicted segmentation mask of CD3/CD8 and the ground-truth mask of the corresponding cells (0 indicates no agreement and 100 indicates perfect agreement). Evaluation of all scores shows that DeepLIIF outperforms all state-of-the-art models. (c) As mentioned earlier, DeepLIIF generalizes across different tissue types and imaging platforms. Two example images from the NuClick Dataset (21) along with the modalities and classified segmentation masks generated by DeepLIIF, are shown in the top rows where the ground-truth mask and quantitative segmentation masks of DeepLIIF and state-of-the-art models are shown in the second row. The mean IOU and Pixel Accuracy are given for each generated mask. (d) Randomly chosen samples from the LYON19 challenge dataset (11). The top row shows the IHC image, and the bottom row shows the classified segmentation mask generated by DeepLIIF. In the mask, the blue color shows the boundary of negative cells, and the red color shows the boundary of positive cells.

We also evaluated the quality of the inferred modalities using mean squared error (MSE) (the average squared difference between the synthetic image and the actual image) and Structural Similarity Index (SSIM) (the similarity between two image). As shown in the Extended Data Figure 2, based on these metrics, DeepLIIF generates highly-realistic images. In this figure, We further visualized the first two components of PCA applied to the feature vectors of synthetic and real images, calculated by the VGG16 model and then applied PCA on the calculated feature vectors. The results show that the synthetic image data points have the same distribution as the real image data points, confirming that the generated images by the model have the same characteristics as the real images. Original/real and DeepLIIF-Inferred modality images of two samples taken from Bladder and Lung tissues are also shown side-by-side with SSIM and MSE scores.

We also tested DeepLIIF on IHC images stained with eight other markers acquired with different scanners and staining protocols. Our testing set includes (1) nine IHC snapshots from a digital microscope stained with Ki67 and PDL1 markers (two examples shown in Extended Data Figure 8), (2) testing set of LYON19 (11) containing 441 IHC CD3/CD8 breast, colon, and prostate ROIs (no annotations) with various staining/tissue artifacts from 8 different institutions (Figure 3(c), and Extended Data Figure 9), (3) PathoNet IHC Ki67 breast cancer dataset (22), containing manual centroid annotations created from consensus of multiple pathologists, acquired in low-resource settings with microscope camera (Extended Data Figure 7), (4) Human Protein Atlas (23) IHC Ki67 (Figure 5) and TP53 images (Extended Data Figure 10), and (5) DLBCL-Morph dataset (24) containing IHC tissue-microarrays for 209 patients stained with BCL2, BCL6, CD10, MYC, MUM1 markers (Extended Data Figure 10.) We visualized the structure of the testing dataset by applying t-distributed stochastic neighbor embedding (t-SNE) to the image styles tested on DeepLIIF in Figure 4. We first extracted the features from each image using the VGG16 model, and applied principal component analysis (PCA) to reduce the number of dimensions in the feature vectors. Next, we visualize the image data points based on the extracted feature vectors using t-SNE. As shown in Figure 4, DeepLIIF is able to adapt to images with various resolutions, color and intensity distributions, and magnifications captured in different clinical settings, and successfully segment and classify the heterogeneous collection of aforementioned testing sets covering eight different IHC markers.

**Figure 4.**
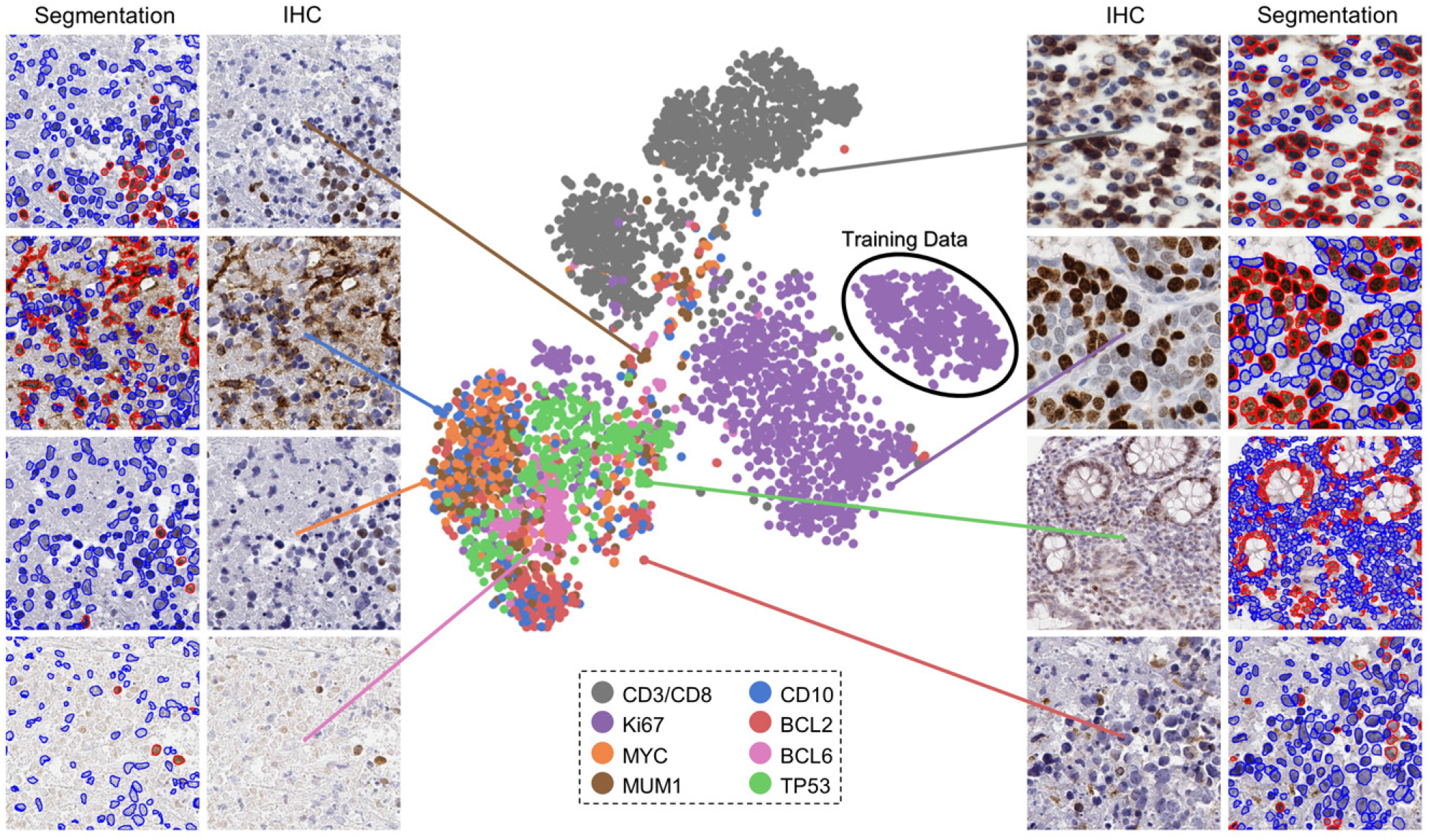
The t-SNE plot of tested IHC markers on DeepLIIF. The structure of the testing dataset is visualized by applying t-SNE to the image styles tested on DeepLIIF. The IHC protein markers in the tested datasets were embedded using t-SNE. Each point represents an IHC image of its corresponding marker. Randomly chosen example images of each marker are shown around the t-SNE plot. The black circle shows the cluster of training images. The distribution of data points shows that DeepLIIF is able to adapt to images with various resolutions, color and intensity distributions, and magnifications captured in different clinical settings, and successfully segment and classify the heterogeneous collection of testing sets covering eight different IHC markers.

**Figure 5.**
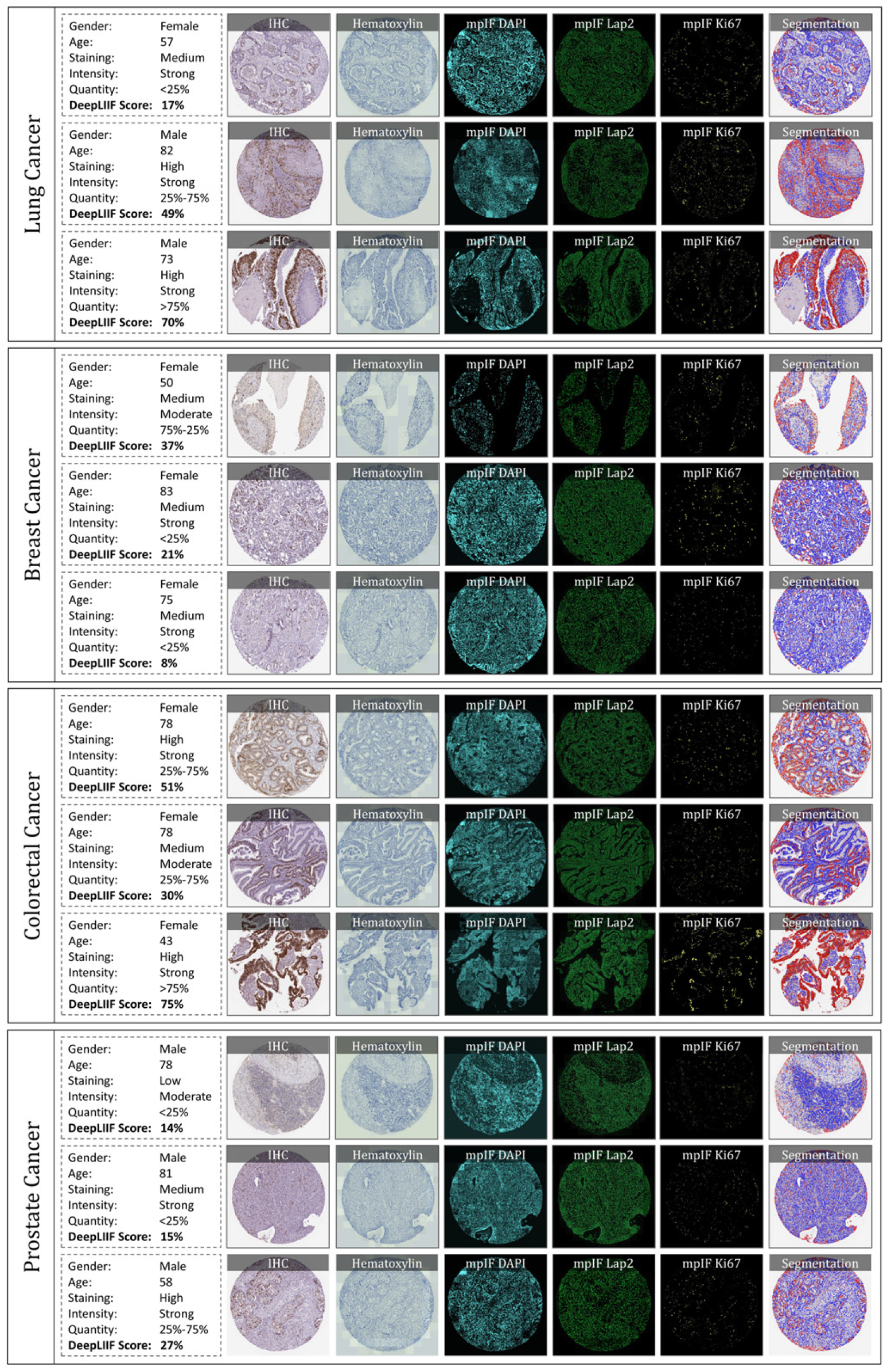
IHC quantification of four cancer type images taken from Protein Atlas IHC Ki67 dataset. In each row, a sample is shown along with the inferred modalities and the classified segmentation mask. The demographic information of the patient and the details about the staining, along with the manual protein score and the predicted score by DeepLIIF are reported next to each sample.

We have also evaluated the performance of DeepLIIF with and without LAP2beta and found the segmentation performance of DeepLIIF with LAP2beta better than without LAP2beta (Extended Data Figure 4). LAP2beta is a nuclear envelope protein broadly expressed in normal tissues. In Extended Data Figure 3, LAP2beta immunohistochemistry reveals nuclear envelope-specific staining in the majority of cells in spleen (99.98%), colon (99.41%), pancreas (99.50%), placenta (76.47%), testis (95.59%), skin (96.74%), lung (98.57%), liver (98.70%), kidney (95.92%) and lymph node (99.86%). Placenta syncytiotrophoblast does not stain with LAP2beta, and the granular layer of skin does not show LAP2beta expression. However, the granular layer of skin lacks nuclei and is therefore not expected to express nuclear envelope proteins. We also observe a lack of consistent Lap2beta staining in the smooth muscle of blood vessel walls (not shown).

DeepLIIF which is solely trained on IHC images stained with Ki67 marker was also tested on H&E images from the MonuSeg Dataset (8). As shown in Extended Data Figure 5, DeepLIIF (out-of-the-box without being trained on H&E images) was able to infer high-quality mpIF modalities and correctly segment the nuclei in these images.

## Discussion

Assessing IHC stained tissue sections is a widely utilized technique in diagnostic pathology laboratories worldwide. IHC-based protein detection in tissue with microscopic visualization is used for many purposes, including tumor identification, tumor classification, cell enumeration, and biomarker detection and quantification. Nearly all IHC stained slides for clinical care are analyzed and reported qualitatively or semi-quantitatively by diagnostic pathologists.

Several approaches have been proposed for deep learningbased stain-to-stain translation of unstained (label-free), H&E, IHC, and multiplex slides, but *relatively few attempts have been made (in limited contexts) at leveraging the translated enriched feature set for cellular-level segmentation, classification or scoring* (25, 26). Recently, Liu et al. (27) used publicly available fluorescence microscopy and histopathology H&E datasets for unsupervised nuclei segmentation in histopathology images by learning from fluorescence microscopy DAPI images. However, their pipeline incorporated CycleGAN, which hallucinated (28) nuclei in the target histopathology domain and hence, required segmentation masks in the source domain to remove any redundant or unnecessary nuclei in the target domain. The model was also not generalizable across the two target histopathology datasets due to the stain variations, making this unsupervised solution less suitable for inferring different cell types from given H&E or IHC images. Burlingame et al. (29) on the other hand, used supervised learning trained on H&E and co-registered single-channel pancytokeratin IF for four pancreatic ductal adenocarcinomas (PDAC) patients to infer pancytokeratin stain for given PDAC H&E image. Another work (30) used a supervised learning method trained on H&E, and co-registered IHC PHH3 DAB slides for mitosis detection in H&E breast cancer WSIs. Recently, Haan et al. (31) used co-registered H&E and special stains for kidney needle core biopsy sections to translate given H&E image to special stains. In essence, there are methods to translate between H&E and IHC but none for translating between IHC and mpIF modalities. *To focus on immediate clinical application, we want to accentuate/disambiguate the cellular information in low-cost IHCs (using a higher-cost and more informative mpIF representation) to improve the interpretability for pathologists as well as for the downstream analysis/algorithms.*

By creating a multitask deep learning framework referred to as DeepLIIF, we provide a unified solution to nuclear segmentation and quantification of IHC stained slides. DeepLIIF is automated and does not require annotations. In contrast, most commercial platforms use a time-intensive workflow for IHC quantification, which involves user-guided (a) IHC-DAB deconvolution, (b) nuclei segmentation of hematoxylin channel, (c) threshold setting for the brown DAB stain, and (d) cell classification based on the threshold. We present a simpler workflow; given an IHC input, we generate different modalities along with the segmented/classified cell masks. Our multitask deep learning framework performs IHC quantification in one process and does not require error-prone IHC deconvolution or manual thresholding steps. We use a single optimizer for all generators and discriminators that improves the performance of all tasks simultaneously. Unique to this model, DeepLIIF is trained by generating registered mpIF, IHC, and hematoxylin staining data from the same slide with the inclusion of nuclear envelope staining to assist in accurate segmentation of adjacent and overlapping nuclei.

Formulating the problem as cell instance segmentation/classification rather than a detection problem helps us to move beyond the reliance on crowd counting algorithms and towards more precise boundary delineation (semantic segmentation) and classification algorithms. DeepLIIF was trained for multi-organ, stain invariant determination of nuclear boundaries and classification of subsequent single-cell nuclei as positive or negative for Ki67 staining detected with the 3,3’-Diaminobenzidine (DAB) chromogen. Subsequently, we determined that DeepLIIF accurately classified all tested nuclear antigens as positive or negative.

Surprisingly, DeepLIIF is often capable of accurate cell classification of non-nuclear staining patterns using CD3, CD8, BCL2, PDL1, and CD10. We believe the success of the DeepLIIF classification of non-nuclear markers is at least in part dependent on the location of the chromogen deposition. BCL2 and CD10 protein staining often show cytoplasmic chromogen deposition close to the nucleus, and CD3 and CD8 most often stain small lymphocytes with scant cytoplasm whereby the chromogen deposition is physically close to the nucleus. DeepLIIF is slightly less accurate in classifying PDL1 staining (Extended Data Figure 8) and, notably, PDL1 staining is more often membranous staining of medium to large cells such as tumor cells and monocyte-derived cell lineages where DAB chromogen deposition is physically further from the nucleus. Since DeepLIIF was not trained for non-nuclear classification, we anticipate that further training using non-nuclear markers will rapidly improve their classification with DeepLIIF.

DeepLIIF, handling of H&E images (Extended Data Figure 5), was the most pleasant surprise where the model out-of-the-box learnt to even separate the H&E images into hematoxylin and (instead of mpIF protein marker) eosin stains. The nuclei segmentations were highly precise. This opens up lot of interesting avenues where we can potentially drive whole slide image registration of neighboring H&E and IHC sections (32) by converting these to a common domain (clean mpIF DAPI images) and then performing deformable image registration.

For IHC images, we have purposely assessed the performance of DeepLIIF for the detection of proteins currently reported semi-quantitatively by pathologists with the goal of facilitating the transition to quantitative reporting if deemed appropriate. We anticipate the further extension of this work to assess the usability of Ki67 quantification in tumors with more unusual morphologic features such as sarcomas. The approach will also be extended to handle more challenging membranous/cytoplasmic markers such as PDL1, Her2, etc as well as H&E and multiplex IHC staining (without requiring any manual/weak annotations for different cell types (3)). Finally, we will incorporate additional mpIF tumor and immune markers into DeepLIIF for more precise phenotypic IHC quantification such as for distinguishing PDL1 expression within tumor vs. macrophage populations.

This work provides a universal, multitask model for both segmenting nuclei in IHC images and recognizing and quantifying positive and negative nuclear staining. Importantly, we describe a modality where training data from higher-cost and higher-dimensional multiplex imaging platforms improves the interpretability of more widely-used and lower-cost IHC.

## Methods

### Training Data

To train DeepLIIF, we used a dataset of lung and bladder tissues containing IHC, hematoxylin, mpIF DAPI, mpIF Lap2, and mpIF Ki67 of the same tissue scanned using ZEISS Axioscan (see Supplementary Materials for more details on staining protocol). These images were scaled and co-registered with the fixed IHC images using affine transformations, resulting in 1667 registered sets of IHC images and the other modalities of size 512 × 512. We randomly selected 709 sets for training, 358 sets for validation, and 600 sets for testing the model.

### Ground-truth Classified Segmentation Mask

To create the ground-truth segmentation mask for training and testing our model, we used our interactive deep learning ImPartial annotations framework (33). Given mpIF DAPI images and few cell annotations, this framework auto-thresholds and performs cell instance segmentation for the entire image. Using this framework, we generated nuclear segmentation masks for each registered set of images with precise cell boundary delineation. Finally, using the mpIF Ki67 images in each set, we classified the segmented cells in the segmentation mask, resulting in 9180 Ki67 positive cells and 59000 Ki67 negative cells. Examples of classified segmentation masks from the ImPartial framework are shown in Figures 1 and 2. The green boundary around the cells are generated by ImPartial, and the cells are classified into red (positive) and blue (negative) using the corresponding mpIF Ki67 image. If a segmented cell has any representation in the mpIF Ki67 image, we classify it as positive (red color), otherwise, we classify it as negative (blue color).

### Objective

Given a dataset of IHC+Ki67 RGB images, our objective is to train a model *f* (.) that maps an input image to four individual modalities, including Hematoxylin channel, mpIF DAPI, mpIF Lap2, and mpIF Ki67 images, and using the mapped representations, generate the segmentation mask. We present a framework, as shown in Figure 6 that performs two tasks simultaneously. First, the translation task translates the IHC+Ki67 image into four different modalities for clinical interpretability as well as for segmentation. Second, a segmentation task generates a single classified segmentation mask from the IHC input and three of the inferred modalities by applying a weighted average and coloring cell boundaries green, positive cells red, and negative cells blue.

**Figure 6.**
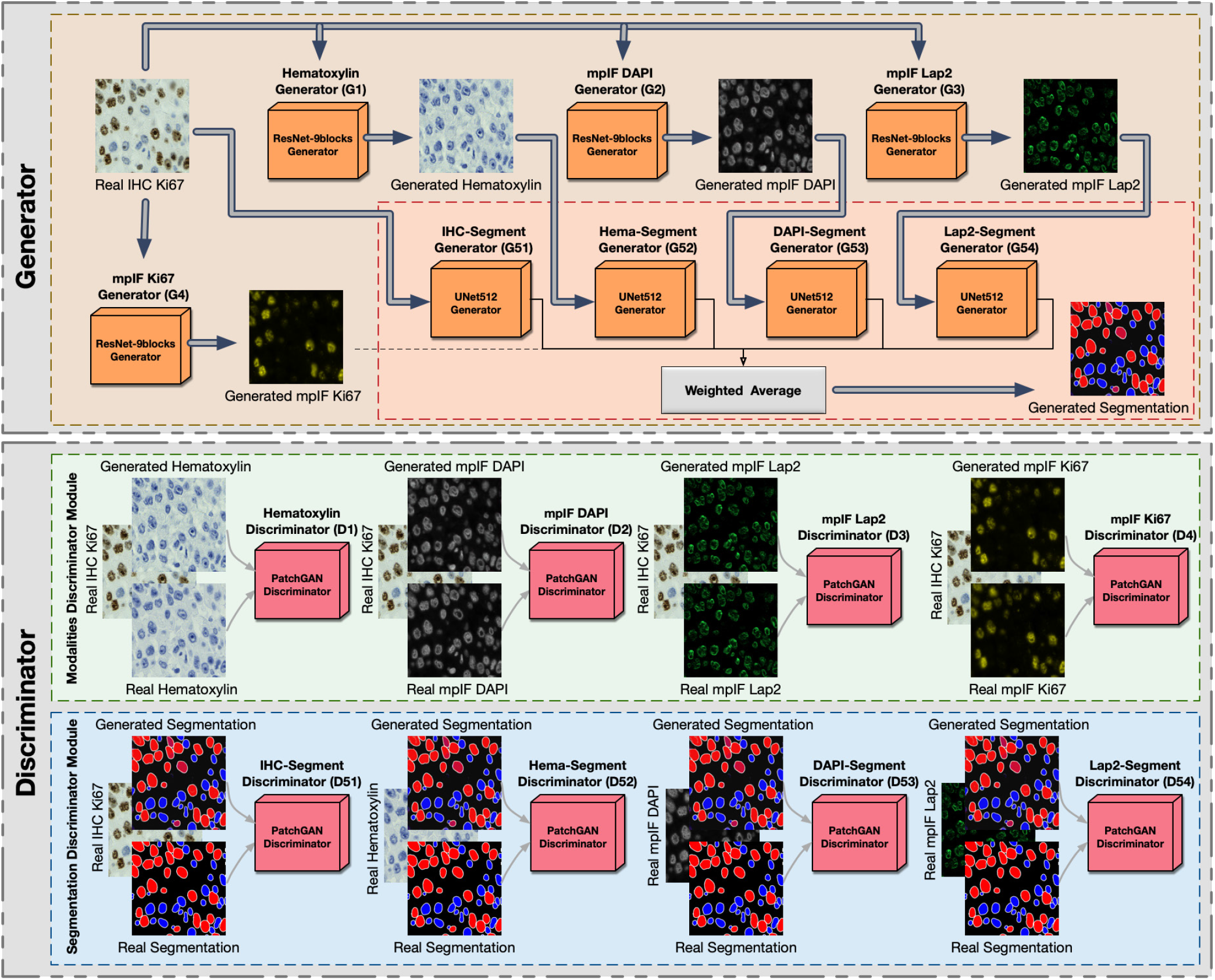
Overview of DeepLIIF. The network consists of a generator and a discriminator component. It uses ResNet-9block generator for generating the modalities including Hematoxylin, mpIF DAPI, mpIF Lap2, and mpIF Ki67 and UNet512 generator for generating the segmentation mask. In the segmentation component, the generated masks from IHC, Hematoxylin, mpIF DAPI, and mpIF Lap2 representations are averaged with pre-defined weights to create the final segmentation mask. The discriminator component consists of the modalities discriminator module and segmentation discriminator module.

We use cGANs to generate the modalities and the segmentation mask. cGANs are made of two distinct components, a generator and a discriminator. The generator learns a mapping from the input image *x* to output image *y, G*: *x* → *y*. The discriminator learns to the paired input and output of the generator from the paired input and ground truth result. We define eight generators to produce four modalities and segmentation masks that cannot be distinguished from real images by eight adversarially trained discriminators (trained to detect fake images from the generators).

### Translation

Generators *G*_*t*_1__, *G*_*t*_2__, *G*_*t*_3__, and *G*_*t*_4__ produce hematoxylin, mpIF DAPI, mpIF Lap2, and mpIF Ki67 images from the input IHC image, respectively (*G_t_i__*: *x_i_* → *y_i_*, where *i* = 1,2,3,4). The discriminator *D_i_* is responsible for discriminating generated images by generators *G_t_i__*. The objective of the conditional GAN for the image translator tasks are defines as follows:

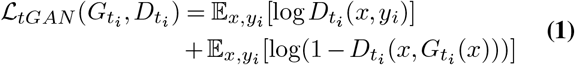

We use smooth L1 loss (Huber loss) to compute the error between the predicted value and the true value, since it is less sensitive to outliers compared to L2 loss and prevents exploding gradients while minimizing blur (34, 35). It is defined as:

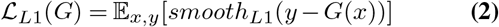

where

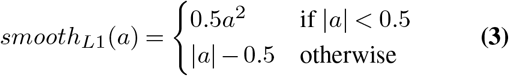

The objective loss function of the translation task is:

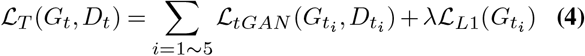

where *λ* controls the relative importance of two objectives.

### Segmentation/Classification

The segmentation component consists of five generators *G*_*S*_1__, *G*_*S*_2__, *G*_*S*_3__, *G*_*S*_4__, and *G*_*S*_5__ producing five individual segmentation masks from the original IHC, inferred hematoxylin image (*G*_*t*_1__), inferred mpIF DAPI (*G*_*t*_2__), inferred mpIF Lap2 (*G*_*t*_3__), and inferred mpIF marker (*G*_*t*_4__), *G*_*S*_*i*__ =: *z_i_* → *y_s_i__* where *i* =1,2,3,4,5. The final segmentation mask is created by averaging the five generated segmentation masks by *G_S_i__* using pre-defined weights, 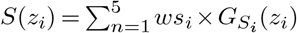, where *ws_i_* are the pre-defined weights. The discriminators *D_S_i__* are responsible for discriminating generated images by generators *G_S_i__*.

In this task, we use LSGAN loss function, since it solves the problem of vanishing gradients for the segmented pixels on the correct side of the decision boundary, but far from the real data, resulting in a more stable boundary segmentation learning process. We define the objective of the conditional GAN for segmentation/classification task as follows:

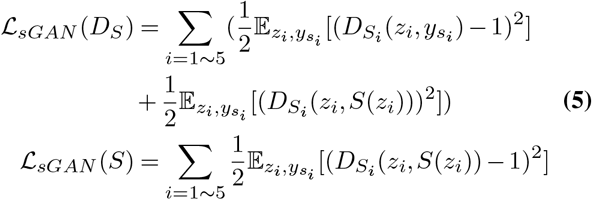

For this task, we also use smooth L1 loss. The objective loss function of the segmentation/classification task is:

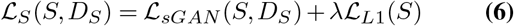

### Final Objective

The final objective is:

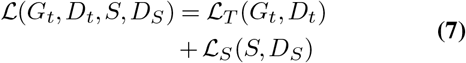

### Generator

We use two different types of generators, ResNet-9blocks generator for producing modalities and U-Net generator for creating segmentation mask.

### ResNet-9blocks Generator

The generators responsible for generating modalities including hematoxylin, mpIF DAPI and mpIF Lap2 starts with a convolution layer and a batch normalization layer followed by Rectified Linear Unit (ReLU) activation function, 2 downsampling layers, 9 residual blocks, 2 upsampling layers, and a covolutional layer followed by a tanh activation function. Each residual block consists of two convolutional layers with the same number of output channels. Each convolutional layer in the residual block is followed by a batch normalization layer and a ReLU activation function. Then, these convolution operations are skipped and the input is directly added before the final ReLU activation function.

### U-Net Generator

For generating the segmentation masks, we use the generator proposed by (35), using the general shape of U-Net (36) with skip connections. The skip connections are added between each layer *i* and layer *n — i* where n is the total number of layers. Each skip connection concatenates all channels at layer *i* with those at layer *n* — *i*.

### Markovian discriminator (PatchGAN)

To address high-frequencies in the image, we use a PatchGAN discriminator that only penalizes structure at the scale of patches. It classifies each *N × N* patch in an image as real or fake. We run this fully convolutional discriminator across the image, averaging all responses to provide the final output of D.

### Optimization

To optimize our network, we use the same standard approach as (37), alternating between one gradient descent step on D and one step on G. In all defined tasks (translation, classification, and segmentation), the network generates different representations for the same cells in the input meaning all tasks have the same endpoint. Therefore, we use a single optimizer for all generators and a single optimizer for all discriminators. Using this approach, optimizing the parameters of a task with a more clear representation of cells improves the accuracy of other tasks since all these tasks are optimized simultaneously.

### Synthetic Data Generation

We found that our model consistently failed in regions with dense clusters of IHC positive cells due to the absence of similar characteristics in our training data. To infuse more information about the clustered positive cells into our model, we developed a novel GAN-based model for the synthetic generation of IHC images using coregistered data. The model takes as input Hematoxylin channel, mpIF DAPI image, and the segmentation mask and generates the corresponding IHC image (Extended Data Figure 6). The model converts the Hematoxylin channel to grayscale to infer more helpful information such as the texture and discard unnecessary information such as color. The Hematoxylin image guides the network to synthesize the background of the IHC image by preserving the shape and texture of the cells and artifacts in the background. The DAPI image assists the network in identifying the location, shape, and texture of the cells to better isolate the cells from the background. The segmentation mask helps the network specify the color of cells based on the type of the cell (positive cell: a brown hue, negative: a blue hue). In the next step, we generated synthetic IHC images with more clustered positive cells. To do so, we change the segmentation mask by choosing a percentage of random negative cells in the segmentation mask (called Neg-to-Pos) and converting these into positive cells. We synthesized new IHC images by setting Neg-to-Pos to 50%, 70%, and 90%. DeepLIIF was retrained with the new dataset, containing original images and these synthesized ones, which resulted in improvement of Dice score by 6.57%, IOU by 7.08%, AJI by 5.53%, and Pixel Accuracy by 2.49%.

### Training Details

We train our model from scratch, using a learning rate of 0.0002 for 100 epochs, and linearly decay the rate to zero over the next 100 epochs. The weights were initialized from a Gaussian distribution N (0, 0.02). We set *λ* = 100 to give more weight to L1 loss. We used batch normalization in our main model. Adam solver (38) was used with a batch size of 1. We use Tree-structured Parzen Estimator (TPE) for hyperparameter optimization, and chose the L1 loss (Least Absolute Deviations) as the evaluation metric to be minimized. We compute the L1 loss for the segmentation mask generated by the model and try to minimize the L1 loss using the TPE approach. We optimized various hyperparameters, including the network generator architecture, the discriminator architecture, the number of layers in the discriminator while using layered architecture, the number of filters in the generator and discriminator, normalization method, initialization method, learning rate, and learning policy, *λ*, and the GAN loss function, segmentation mask generators weights with diverse options for each of them. Based on the hyperparameter optimization, the following predefined weights (*ws_i_*) were set for individual modalities to generate the final segmentation mask: weight of segmentation mask generated by original IHC image (*ws*_1_)= 0.25, Hematoxylin channel (*ws*_2_)= 0.15, mpIF DAPI (*ws*_3_) = 0.25, mpIF Lap2 (*ws*_4_)= 0.1, and mpIF protein marker image (*ws*_5_)= 0.25. The cell type (positive or negative) is classified using the original IHC image (where brown cells are positive and blue cells are negative) and the mpIF protein marker image (which only shows the positive cells). Therefore, to have enough information on the cell types, these two representations are assigned 50% of the total weight with equal contribution. The mpIF DAPI image contains the representation of the cell where the background and artifacts are removed. Since this representation has the most useful information on the cell shape, area, and boundaries, it was assigned 25% of the total weight in creating the segmentation mask. The mpIF Lap2 image is generated from the mpIF DAPI image and it contains only the boundaries on the cells. Even though it has more than 90% coverage, it still misses out on cells, hence 15% of the total weight makes sense. With this weightage, we can be sure that if there is any confusing information in the mpIF DAPI image, it does not get infused into the model by a large weight. Also, by giving less weight to the Lap2, we increase the final segmentation probability of the cells not covered by Lap2. The Hematoxylin image has all the information, including the cells with lower intensities, the artifacts, and the background. Since this image shares the background and artifacts information with the IHC image and the cell information with the mpIF DAPI image, it is given less weight to decrease the probability of artifacts being segmented and classified as cells.

One of the challenges in GANs is the instability of its training. We used spectral normalization, a weight normalization technique, to stabilize the training of the discriminator (39). Spectral normalization stabilizes the training of discriminators in GANs by re-scaling the weight tensor with spectral norm *σ* of the weight matrix calculated using the power iteration method. If the dimension of the weight tensor is greater than 2, it is reshaped to 2D in the power iteration method to get the spectral norm. We first trained the model using spectral normalization on the original dataset. The spectral normalization could not significantly improve the performance of the model. The original model achieved Dice score of 61.57%, IOU 46.12%, AJI 47.95% and Pixel Accuracy 91.69% whereas the model with spectral normalization achieved a Dice score of 61.57%, IOU of 46.17%, AJI of 48.11% and Pixel Accuracy of 92.09%. In another experiment, we trained the model with spectral normalization on our new dataset containing original as well as the generated synthetic IHC images. The Dice score, IOU, and Pixel accuracy of the model trained using spectral normalization dropped from 68.15% to 65.14%, 53.20% to 51.15%, and 94.20% to 94.18%, respectively, while the AJI improved from 53.48% to 56.49%. As the results show, the addition of the synthetic images in training improved the model’s performance across all metrics.

To increase the inference speed of the model, we also experimented with many-to-one approach for segmentation/classification task to decrease the number of generators to one. In this approach, we still have four generators and four discriminators for inferring the modalities but use one generator and one discriminator (instead of five) for segmentation/classification task, trained on the combination of all inferred modalities. We first trained this model with the original dataset. Compared to the original model with five segmentation generators, the Dice score, IOU, AJI, and Pixel Accuracy dropped by 12.13%, 10.21%, 12.45%, and 3.66%, respectively. In another experiment, we trained the model with one segmentation generator on the new dataset including synthetic images. Similar to the previous experiment, using one generator instead of five independent generators deteriorated the model’s performance in terms of Dice score by 7%, IOU by 6.49%, AJI by 3.58%, and Pixel Accuracy by 0.98%. We observed that similar to the original model, the addition of synthetic IHC images in the training process with one generator could increase the Dice score from 49.44% to 61.13%, the IOU from 35.91% to 46.71%, the AJI from 35.50% to 49.90%, and Pixel Accuracy from 88.03 to 93.22%, while reducing the performance drop, compared to the original model; this was still significantly less than the best performance from the multi-generator configuration, as shown above, Dice score 68.15%, IOU 53.20%, AJI 53.48%, and Pixel Accuracy 94.20%.

### Testing Details

The inference time of the model for a patch of 512 × 512 is 4 seconds. To infer modalities and segment an image larger than 512 × 512, we tile the image into overlapping patches. The tile size and overlap size can be given by the user as an input to the framework. The patches containing no cells are ignored in this step, improving the inference time. Then, we run the tiles through our model. The model resizes the given patches to 512 for inference. In the final step, we stitch tiles using the given overlap size to create the final inferred modalities and the classified segmentation mask. It takes about 10 to 25 minutes (depending on the percentage of cell-containing region, the WSI magnification level, user-selected tile size and overlap size) to infer the modalities and the classified segmentation mask of a WSI with size of 10000 × 10000 with 40x magnification.

### Ablation Study

DeepLIIF infers four modalities to compute the segmentation/classification mask of an IHC image. We perform an ablation study on each of these four components. The goal of this experiment is to investigate if the performance improvements are due to the increased ability of each task-specific network to share their respective features. In each experiment, we trained our model with three modalities, each time removing a modality to study the accuracy of the model in absence of that modality. We tested all models on the BC Dataset of 164 images with size 512 × 512. The results show that the original model (with all modalities) with Dice score 65.14%, IOU 51.15%, AJI 56.49% and Pixel Accuracy of 94.20% outperforms the model without Hematoxylin modality with Dice score 62.86%, IOU 47.68%, AJI 50.10% and Pixel Accuracy 92.43%, model without mpIF DAPI with Dice score 62.45%, IOU 47.13%, AJI 50.38% and Pixel Accuracy 92.35%, model without mpIF Lap2 with Dice score 61.07%, IOU 45.71%, AJI 49.14%, and Pixel Accuracy 92.16%, and model without mpIF protein marker with Dice score 57.92%, IOU 42.91%, AJI 47.56%, and Pixel Accuracy 91.81%. The mpIF Lap2 is important for splitting overlapping cells and detecting boundaries (the model without mpIF Lap2 has the lowest AJI score). Moreover, mpIF Lap2 is the only modality among the four that clearly outlines the cells in regions with artifacts or noise. The model without mpIF protein marker image has the worst Pixel Accuracy and Dice score, showing its clear importance in cell classification. The mpIF DAPI image guides the model in predicting the location of the cells, given the drop in Pixel Accuracy and AJI score. Hematoxylin image on the other hand seems to make the least difference when removed, though it helps visually (according to two trained pathologists) by providing a separated hematoxylin channel from the IHC (Hematoxylin + DAB) input.

## Supporting information

Supplement_staining_protocol

## Data Availability

The complete IHC Ki67 BCDataset with manual annotations is available at https://sites.google.com/view/bcdataset. Complete lymphocytes detection IHC CD3/CD8 (LYON challenge) dataset is available at https://zenodo.org/record/3385420#.XW-6JygzYuW. The NuClick IHC annotations for crops from the LYON19 dataset can be found at https://warwick.ac.uk/fac/sci/dcs/research/tia/data/nuclick/ihc_nuclick.zip. DLBCL-Morph dataset with BCL2, BCL6, MUM1, MYC, and CD10 IHCs is accessible at https://stanfordmedicine.box.com/s/ub8e0wlhsdenyhdsuuzp6zhj0i82xrb1.

The high-res tiff images for TP53 IHCs can be downloaded from https://www.proteinatlas.org/ENSG00000141510-TP53. All our internal training and testing data (acquired under the IRB protcol approval #16-1683), and source data underlying figures (in excel files) along with the pretrained models are available at https://zenodo.org/record/4751737#.YV379XVKhH4.

## Code Availability

All code was implemented in Python using PyTorch as the primary deep learning package. All code and scripts to reproduce the experiments of this paper are available at https://github.com/nadeemlab/DeepLIIF and releases are available at https://doi.org/10.5281/zenodo.5553268. For convenience, we have also included docker file as well as Google CoLab Demo project (in case someone does not have access to a GPU and wants to run their images directly via the CoLab project). The Google CoLab project can be accessed at https://colab.research.google.com/drive/12zFfL7rDAtXfzBwArh9hb0jvA38L_ODK?usp=sharing.

## Acknowledgements

This project was supported by MSK Cancer Center Support Grant/Core Grant (P30 CA008748) and in part by MSK Dig-ITs Hybrid Research Initiative and NSF grants CNS1650499, OAC1919752, and ICER1940302.

## Author Contributions

SN, TJH, and PG conceived the study and designed the experiments. SN and PG wrote the computer codes and performed the experimental analysis. YL and TJH performed the IHC and multiplex staining. MA, TJH, and NG conceived the Lap2BETA idea for nuclear envelop staining. PG, SN, AK, and RV analyzed the results. SN, TJH and PG prepared the manuscript with input from all co-authors. SN supervised the research.

## Competing Interests

The authors declare no competing interests.

---

**Extended Data Figure 1.**
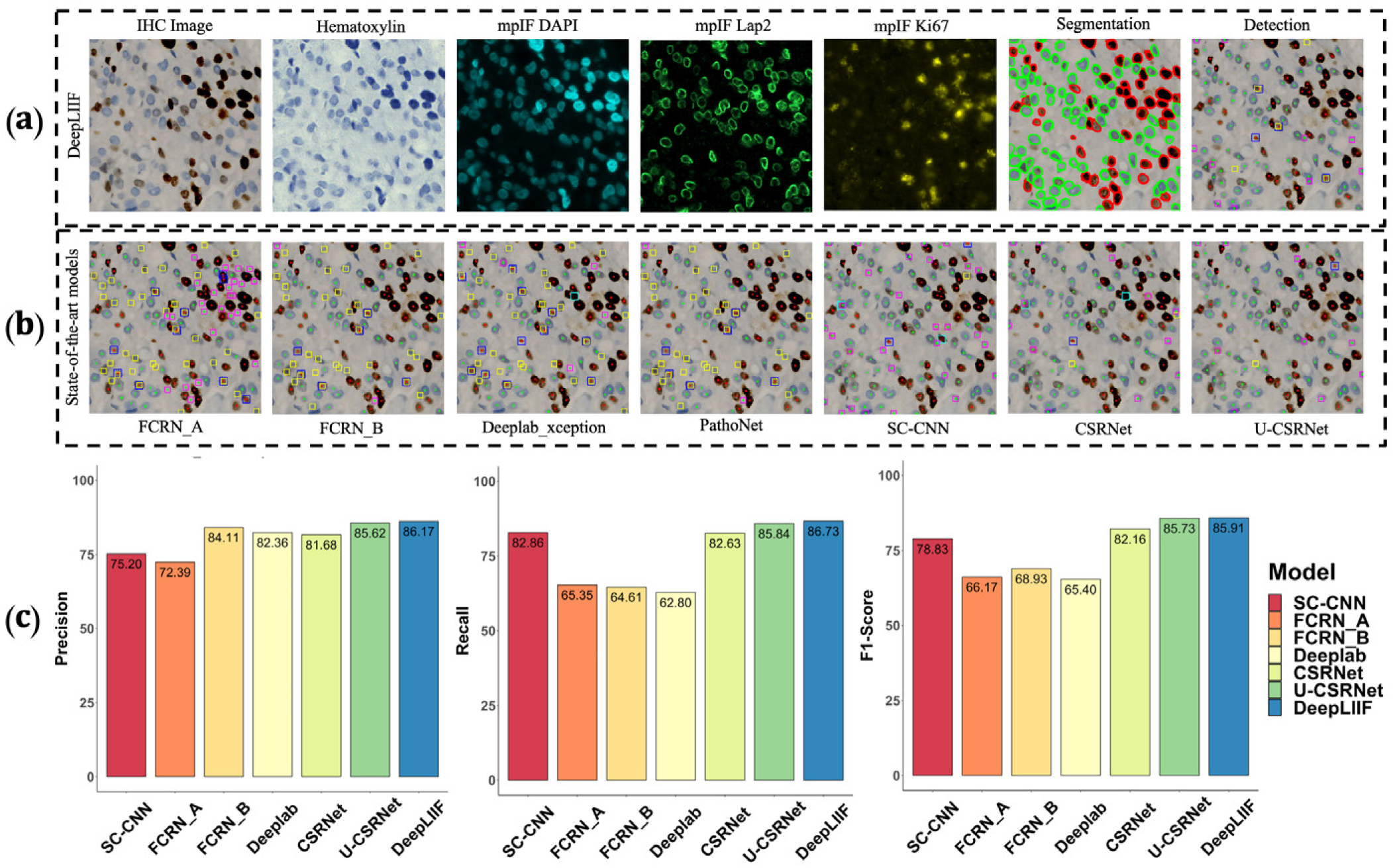
Qualitative and quantitative analysis of DeepLIIF against detection models on the testing set of the BC Data (9). (a) An example IHC image from the BC Data testing set, the generated modalities, segmentation mask overlaid on the IHC image, and the detection mask generated by DeepLIIF (b) The detection masks generated by the detection models. In the detection mask, the center of a detected positive cell is shown with red dot and the center of a detected negative cell is shown with blue dot. We show the missing positive cells in cyan bounding boxes, the missing negative cells in yellow bounding boxes, the wrongly detected positive cells in blue bounding boxes, the wrongly detected negative cells in pink bounding boxes. (c) The detection accuracy is measured by getting average of precision 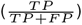, recall 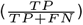, and f1-score 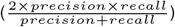 between the predicted detection mask of each class and the ground-truth mask of the corresponding class. A predicted point is regarded as true positive if it is within the region of a ground-truth point with a predefined radius (we set it to 10 pixels in our experiment which is similar to the predefined radius in (9)). Centers that have been detected more than once are considered as false positive. Evaluation of all scores show that DeepLIIF outperforms all state-of-the-art models.

**Extended Data Figure 2.**
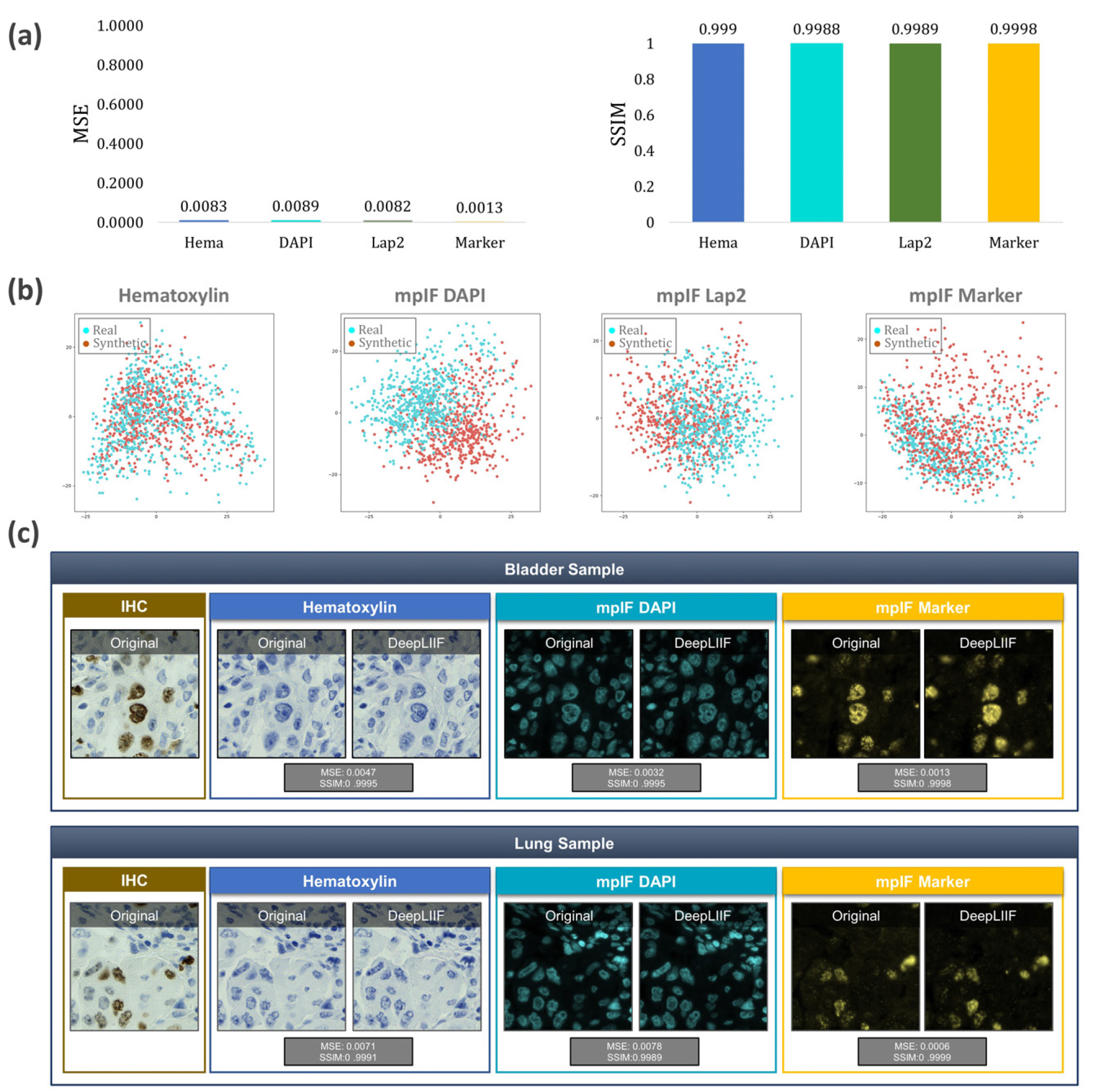
Quantitative and qualitative analysis of DeepLIIF on modality inference. (a) The Quantitative analysis of the synthetic data against the real data using MSE, SSIM, Inception Score, and FID. The low value of MSE (close to 0) and the high value of SSIM (close to 1) shows that the model generates high quality synthetic images similar to real images. (b) Visualization of first two components of PCA applied to synthetic and real images. We first, calculated a feature vector for each image using VGG16 model and then we applied PCA on the calculated feature vectors and visualized the first two components. As shown in the figure, the synthetic image data points have the same distribution as the real image data points, showing that the generated images by the model have the same characteristics as the real images. (c) The original/real and model-inferred modalities of two samples taken from Bladder and Lung tissues are shown side-by-side.

**Extended Data Figure 3.**
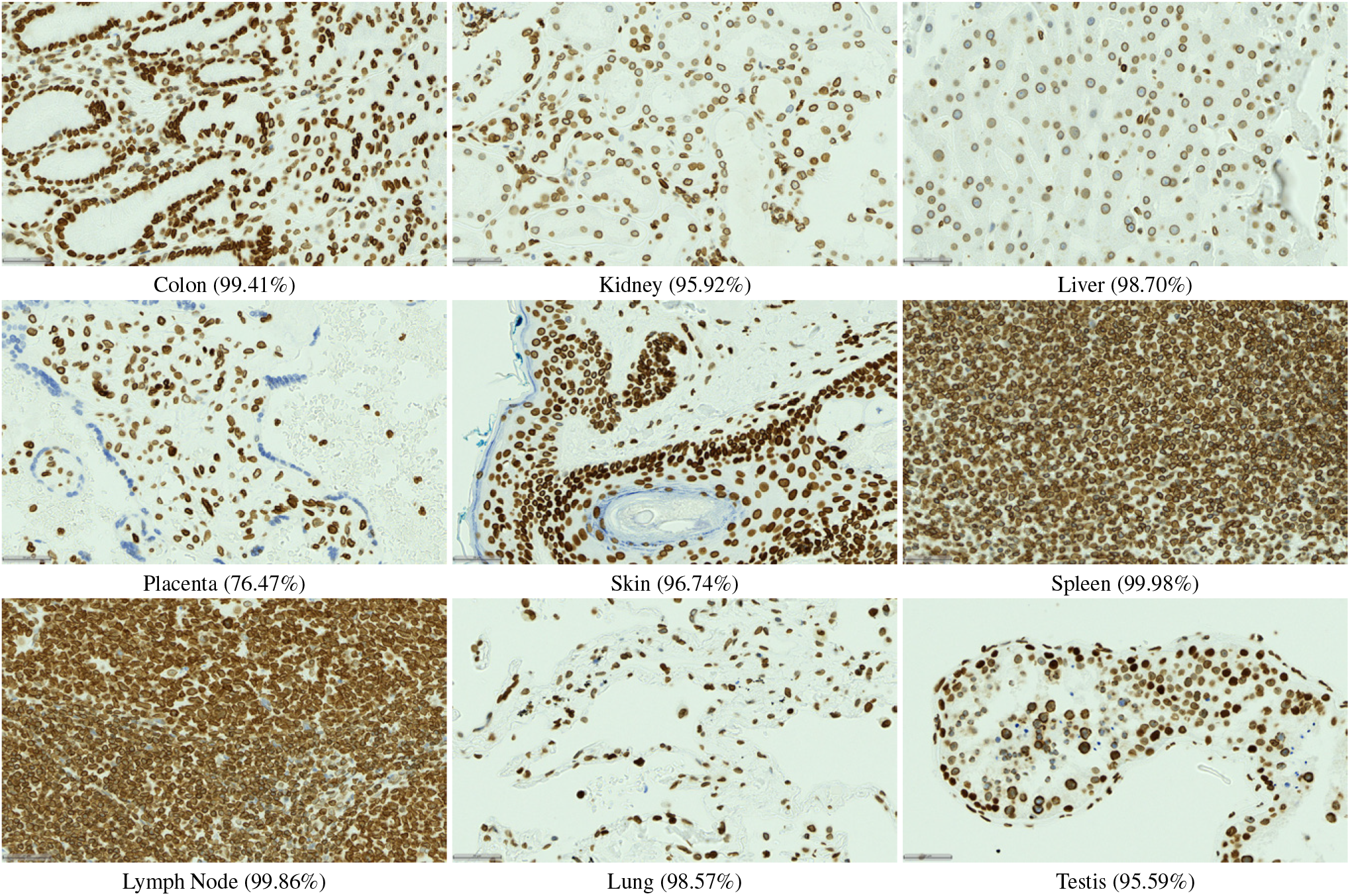
LAP2beta coverage for normal tissues. LAP2beta immunohistochemistry reveals nuclear envelope-specific staining in the majority of cells in spleen (99.98%), colon (99.41%), pancreas (99.50%), placenta (76.47%), testis (95.59%), skin (96.74%), lung (98.57%), liver (98.70%), kidney (95.92%) and lymph node (99.86%).

**Extended Data Figure 4.**
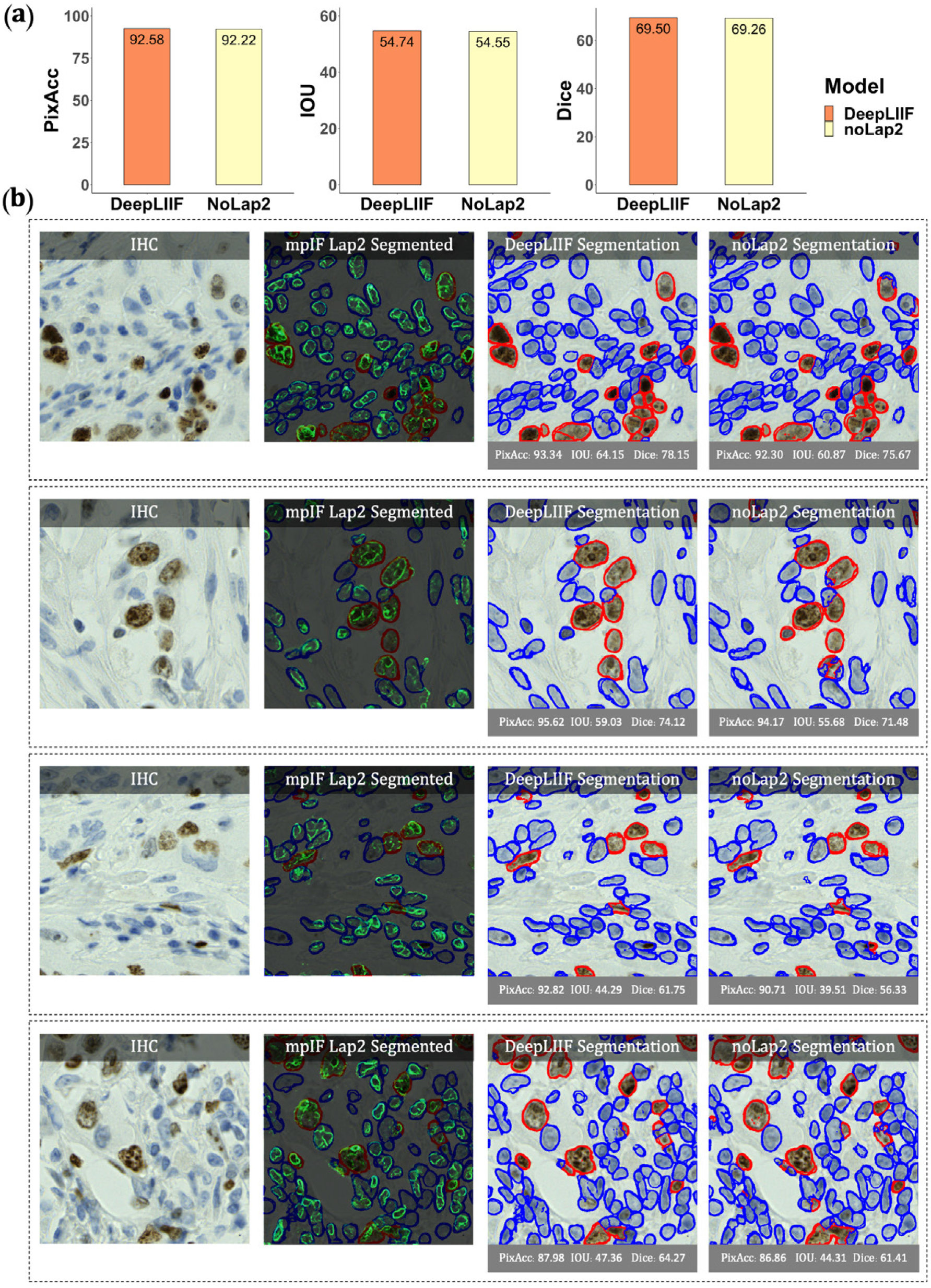
Qualitative and quantitative analysis of DeepLIIF against the same model without using mpIF Lap2, referred to as noLap2 model. (a) A qualitative comparison of DeepLIIF against noLap2 model. (b) Some example IHC images. The first image in each row shows the input IHC image. In the second image, the generated mpIF Lap2 image is overlaid on the classified/segmented IHC image. The third and fourth images show the segmentation mask, respectively, generated by DeepLIIF and noLap2.

**Extended Data Figure 5.**
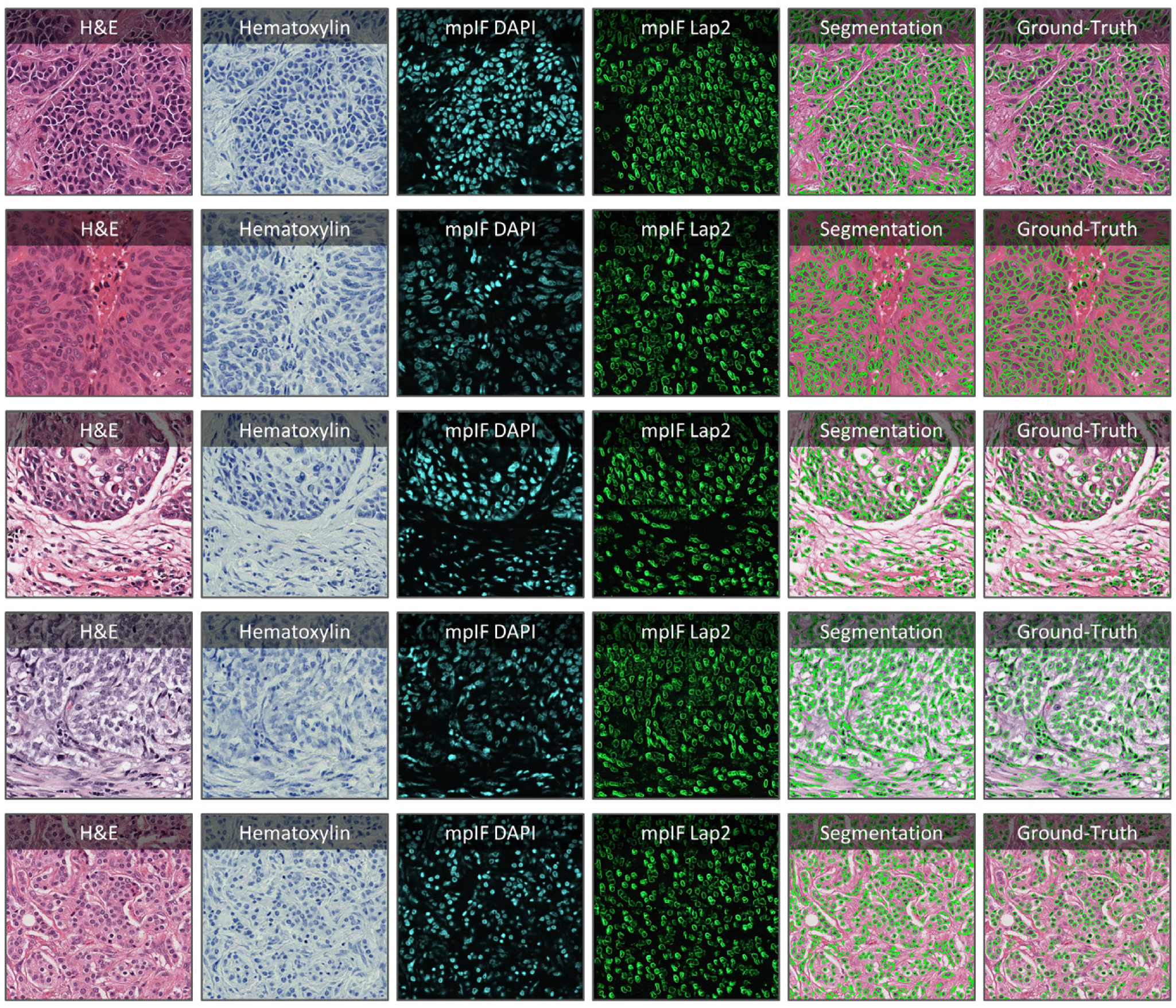
Application of DeepLIIF on some H&E sample images taken from MonuSeg Dataset (8). We tested DeepLIIF, trained solely on IHC images stained with Ki67 marker, on H&E images. In each row, the inferred modalities and the segmentation mask overlaid on the original H&E sample are shown.

**Extended Data Figure 6.**
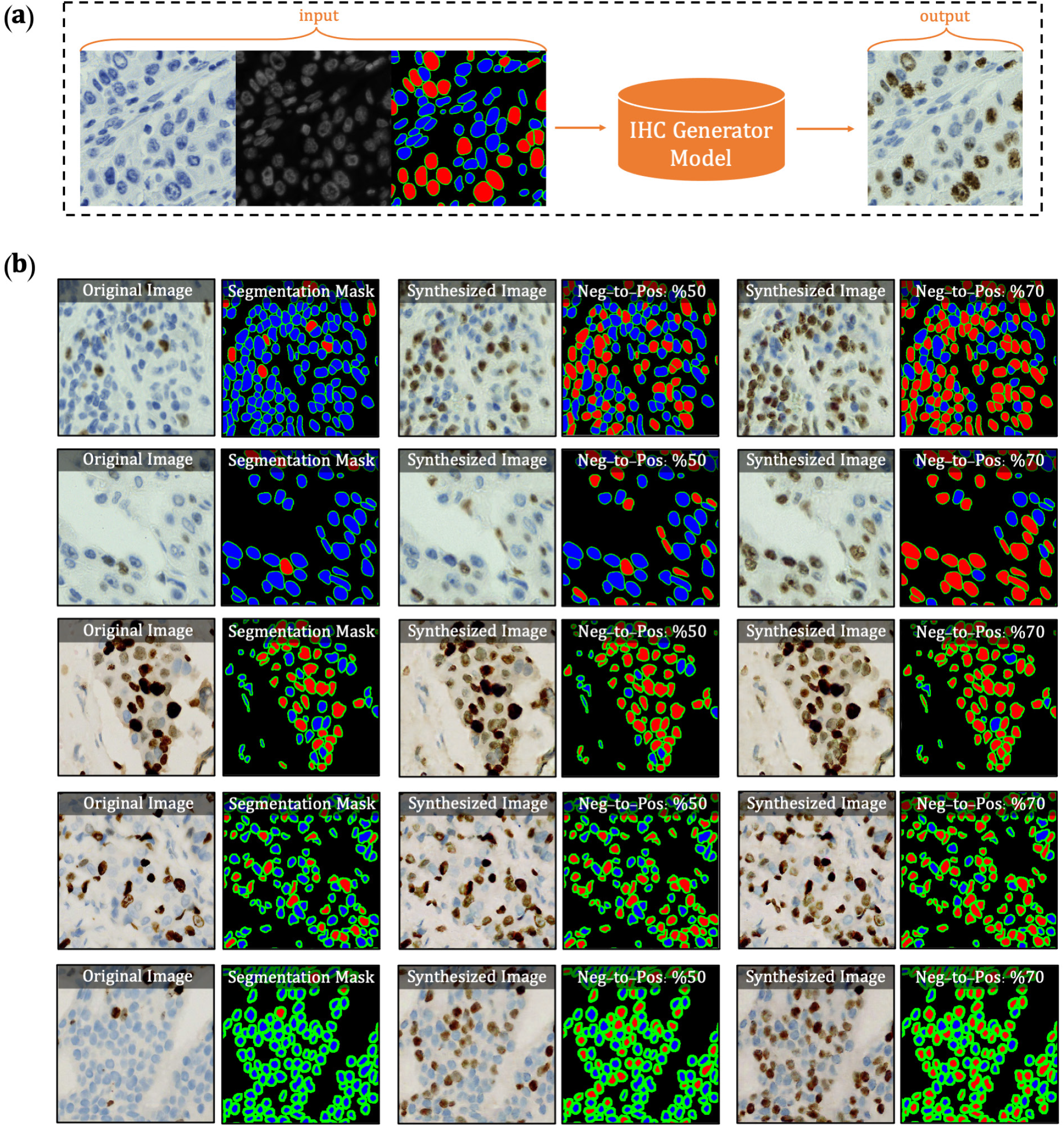
Overview of synthetic IHC image generation. (a) A training sample of the IHC-generator model. (b) Some samples of synthesized IHC images using the trained IHC-Generator model. The Neg-to-Pos shows the percentage of the negative cells in the segmentation mask converted to positive cells.

**Extended Data Figure 7.**
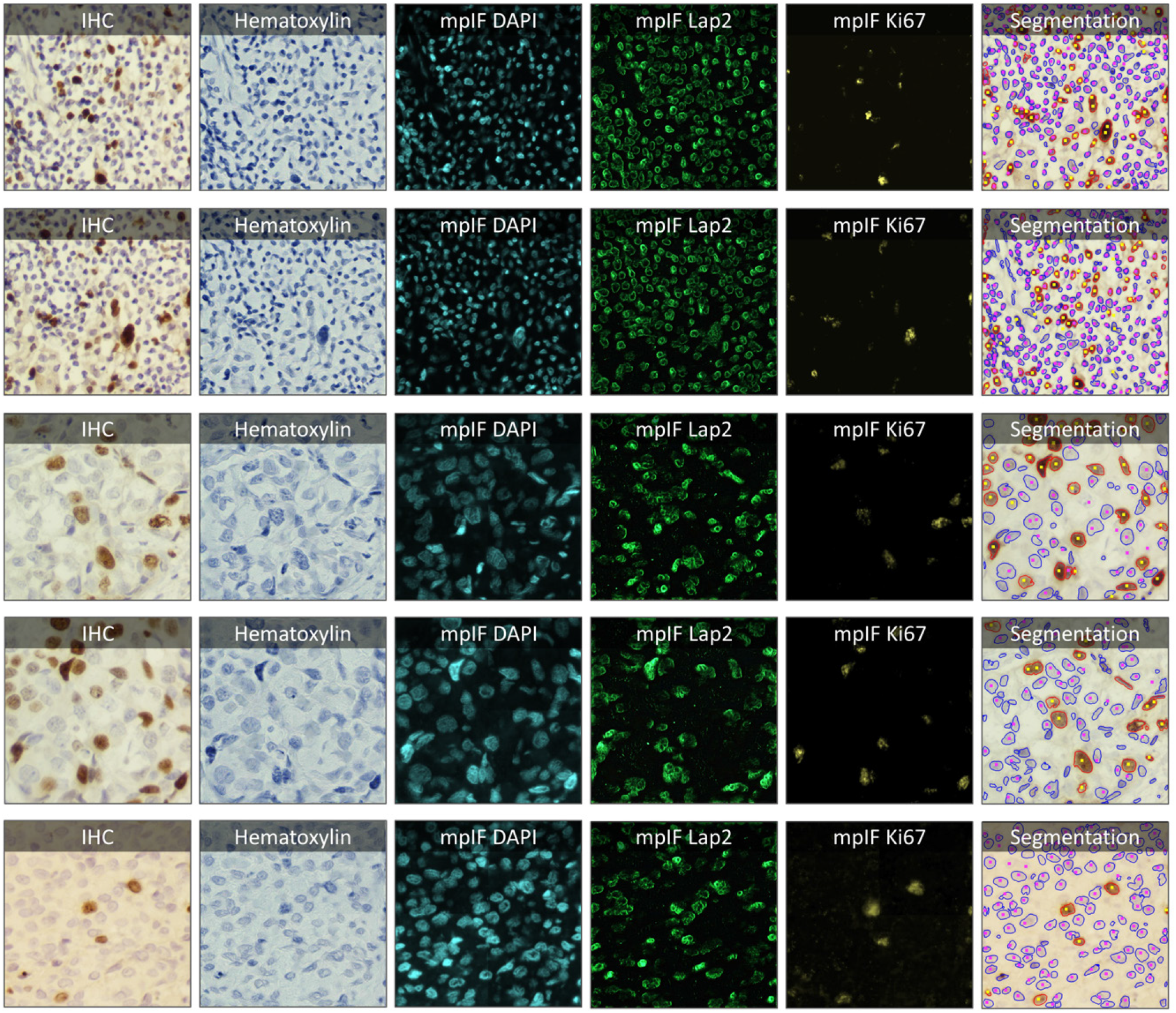
Samples taken from the PathoNet IHC Ki67 breast cancer dataset (22) along with the inferred modalities and classified segmentation mask marked by manual centroid annotations created from consensus of multiple pathologists. The IHC images were acquired in low-resource settings with microscope camera. In each row, the sample IHC image along with the inferred modalities are shown. The overlaid classified segmentation mask generated by DeepLIIF with manual annotations are shown in the furthest right column. The blue and red boundaries represent the negative and positive cells predicted by the model, while the pink and yellow dots show the manual annotations of the negative and positive cells, respectively.

**Extended Data Figure 8.**
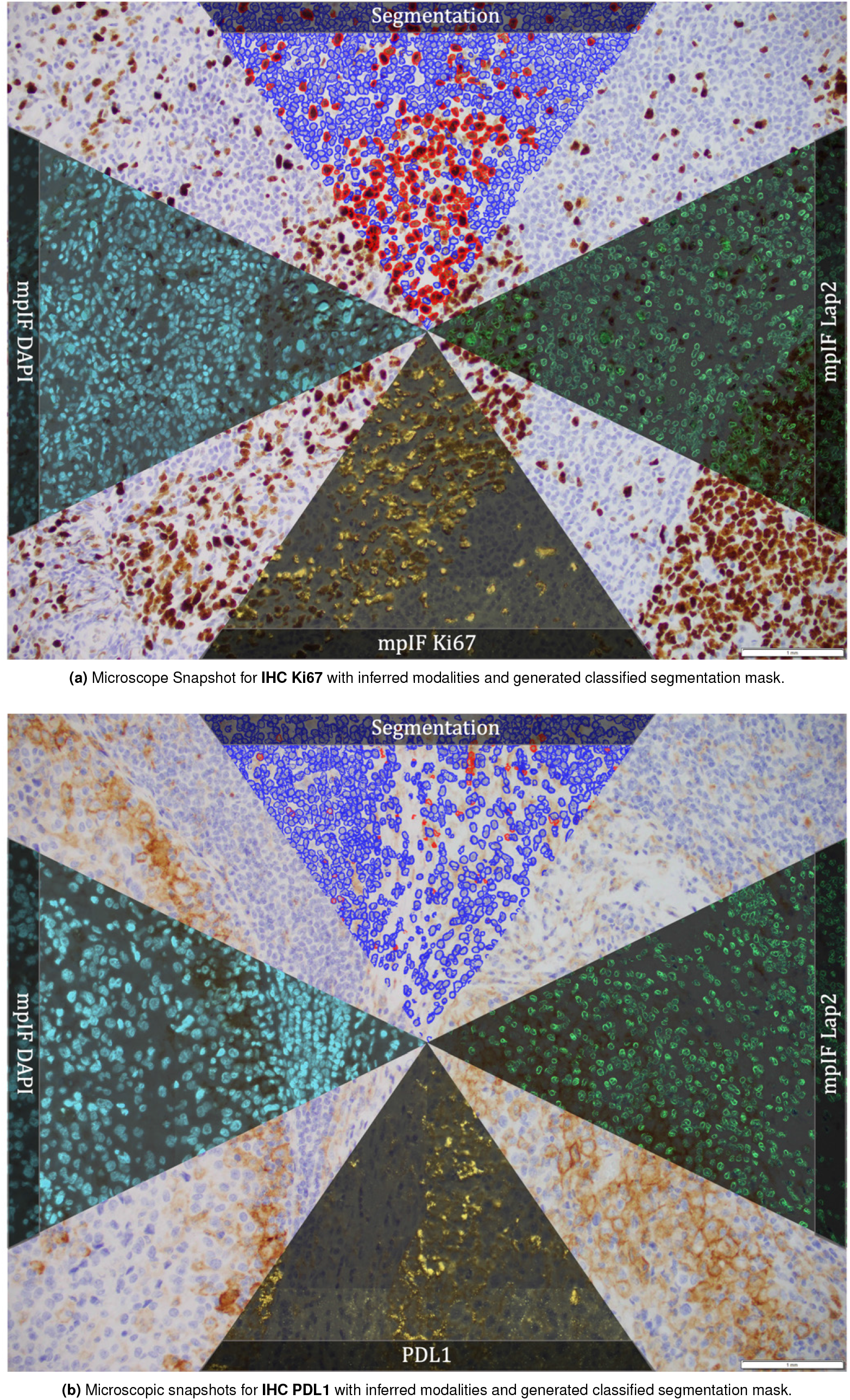
Microscopic snapshots of IHC images stained with two different markers along with inferred modalities and generated classified segmentation mask.

**Extended Data Figure 9.**
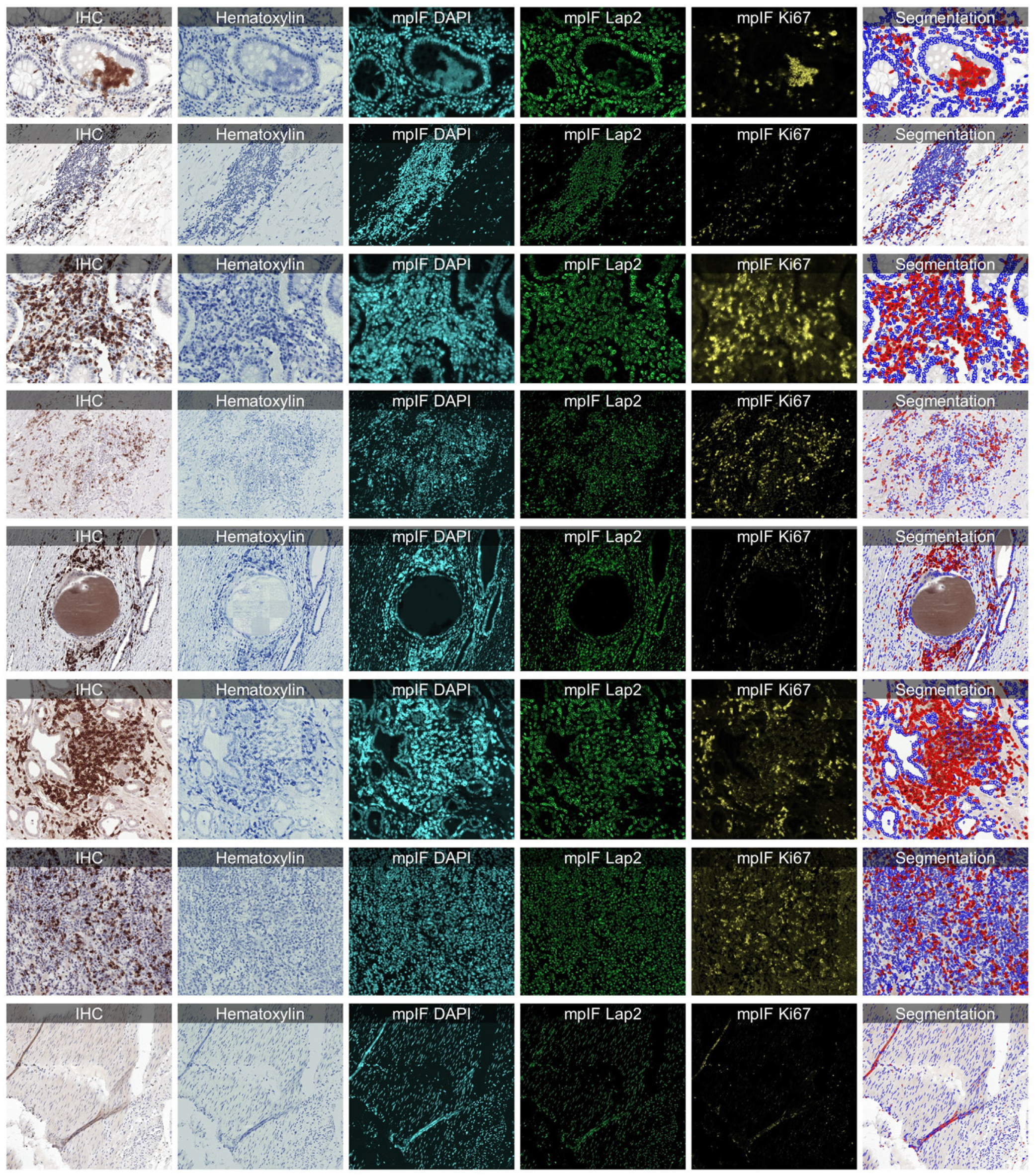
Some examples from LYON19 Challenge Dataset (11). The generated modalities and classified segmentation mask for each sample are in a separate row.

**Extended Data Figure 10.**
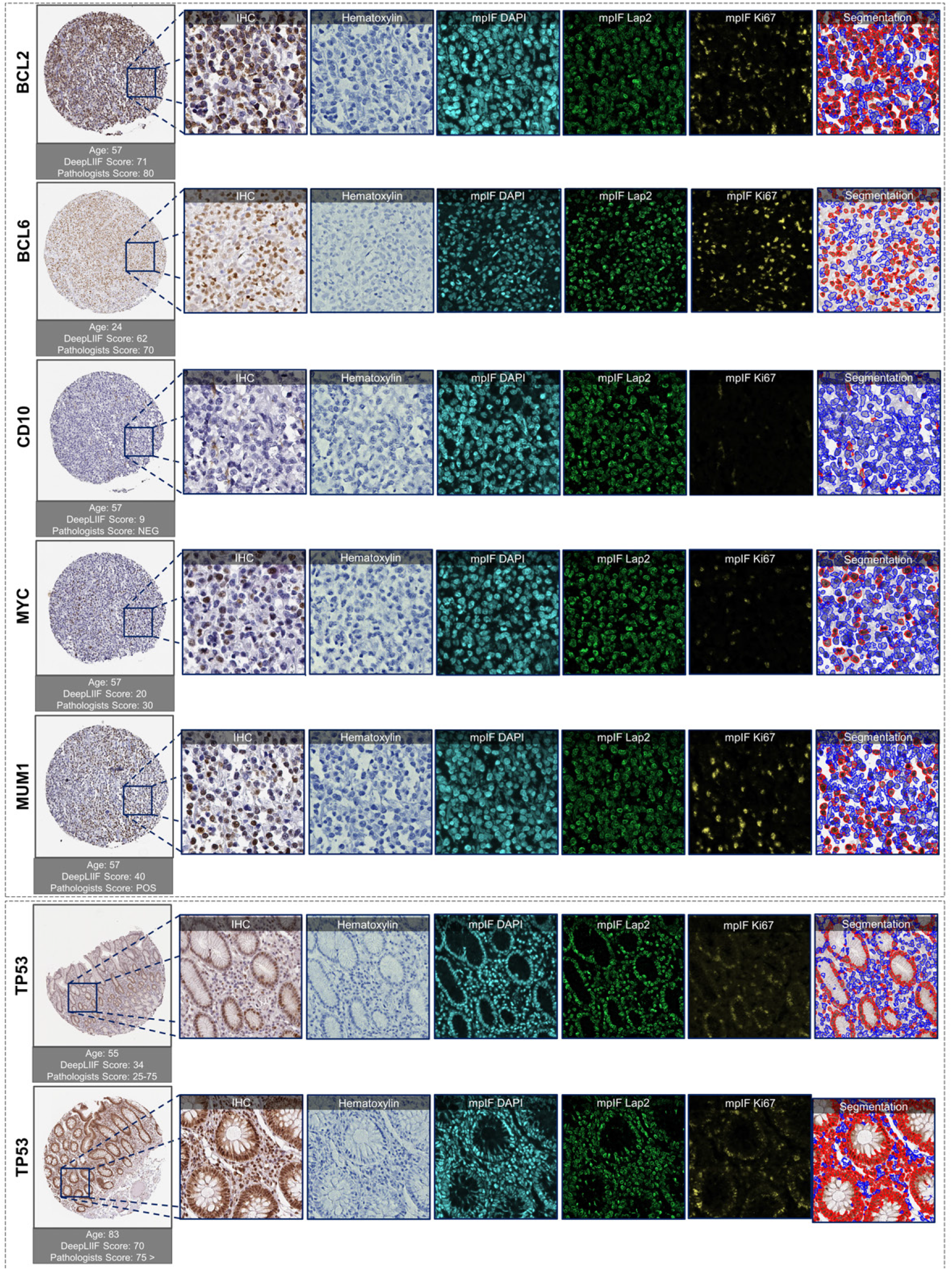
Examples of tissues stained with various markers. The top box shows sample tissues stained with BCL2, BCL6, CD10, MYC, and MUM1 from DLBCL-morph dataset (24). The bottom box shows sample images stained with TP53 marker from the Human Protein Atlas (23). In each row, the first image on the left shows the original tissue stained with a specific marker. The quantification score computed by the classified segmentation mask generated by DeepLIIF is shown on the top of the whole tissue image, and the predicted score by pathologists is shown on the bottom. In the following images of each row, the modalities and the classified segmentation mask of a chosen crop from the original tissue are shown.

## Notes

https://github.com/nadeemlab/DeepLIIF

### Competing Interest Statement

The authors have declared no competing interest.

### Summary of Updates

Revised manuscript

