## Supplement_staining_protocol for "Deep Learning-Inferred Multiplex ImmunoFluorescence for IHC Image Quantification"

### Tissue Staining and Imaging

**First round: Hoechst single staining and imaging**

4 µm FFPE tissue sections sectioned to "superfrost plus microscope slides" (Fisher Scientific 12-550-15) were baked for 3 hrs. at 62 degrees Celsius in vertical slide orientation with subsequent prestaining with deparaffinization and 30 minutes of antigen retrieval with Leica Bond ER2 was performed on the Leica Bond RX followed by Hoechst (Invitrogen 33342) nuclei staining and mounted with ProLong Gold antifade reagent mounting medium (Invitrogen P36930). The full slide imaging was acquired with Zeiss AXIO Scanner at 20X (200X final magnification). The DAPI filter cube was used for the imaging (Semrock Brighline LED-DAPI-B-ZHE-ZERO).

**Second round: 3-Color Multiplex immunoflourescence staining and imaging**

The DAPI stained slide was passively de-coverslipped by soaking in PBS until the coverslip falls off and then the slide was washed with PBS to remove the mounting medium and multiplex IF staining and imaging with the following method:

Primary antibody staining conditions were optimized using standard immunohistochemical staining on the Leica Bond RX automated research stainer with DAB detection (Leica Bond Polymer Refine Detection DS9800). Using 4 µm formalin-fixed, paraffin-embedded tissue sections and serial antibody titrations, the optimal antibody concentration was determined followed by transition to multiplex assay with equivalency. Optimal primary antibody stripping conditions between rounds in the 3-color assay were performed following 1 cycle of tyramide deposition followed by heat-induced stripping (see below) and subsequent chromogenic development (Leica Bond Polymer Regine Detection DS9800) with visual inspection for chromogenic product with a light microscope (TH). Multiplex assay antibodies and conditions are described in Table 1.

**Table 1**.

| Antigen | Antibody Clone | Manufacturer | Titration | Detection Dye (cycle) |
|---|---|---|---|---|
| Lap2-Beta | 27/LAP2 | BD | 0.33µg/ml | Opal 520 (1) |
| Ki67 | SP6 | Biocare | 1:100 | Opal 570 (2) |
| panCK | AE1/AE3 | DAKO | 0.665µg/ml | Opal 650 (3) |

**Three-color multiplex imaging assay:**

Three sequential cycles of staining with each round including a 10-minute blocking (Akoya antibody diluent/block ARD1001) and a 30-minute primary antibody incubation. Detection of primary antibodies was performed using a goat anti-mouse Poly HRP secondary antibody (Lap2-beta and panCK) or goat anti-rabbit Poly HRP secondary antibody (Ki67; IInvitrogen B40961/2; 10-minute incubation). The HRP-conjugated secondary antibody polymer was detected using fluorescent tyramide signal amplification using Opal dyes 520, 570, 650(Akoya FP1487001KT, FP1488001KT, FP1496001KT). The covalent tyramide reaction was followed by heat induced stripping of the primary/secondary antibody complex using Akoya AR9 buffer (AR900250ML) and Leica Bond ER2 (90% ER2 and 10% AR9) at 100 degrees Celsius for 20 minutes preceding the next cycle (1 cycle of stripping for Lap2-Beta, panCK, two cycles for Ki67). After 3 sequential rounds of staining, sections were stained with Hoechst (Invitrogen 33342) to visualize nuclei and mounted with ProLong Gold antifade reagent mounting medium (Invitrogen P36930).

Three color multiplex stained slides were imaged using the Zeiss AXIO Scanner. Full slide imaging was acquired at 20X (200X final magnification). Filter cubes used for multispectral imaging were DAPI, FITC, Cy3, and Cy5.

**Third round: Hematoxylin single staining and imaging** The multiplex IF slide(s) was de-coverslipped as previously described and slides were washed with PBS to remove mounting medium. Hematoxylin staining was performed on the Leica BondRX. Slides were mounted with EcoMount (Biocare, EM897L) with addition of a coverslip (Corning 2975-224) with subsequent bright field full slide imaging with Zeiss AXIO Scanner.

**Fourth round: Repeat the de-coverslip step with standard immunohistochemical staining of Ki67**(SP6, Biocare) on the Leica Bond RX automated research stainer using DAB detection (Leica Bond Polymer Refine Detection DS9800). Standard tissue dehydration processing with gradient Ethanol (Sigma, R8382-1GA) and xylene (epredia, 6601) was used and slides were mounted with EcoMount (Biocare, EM897L). Full slide imaging was acquired at 20X (200X final magnification) with Zeiss AXIO Scanner.

**Fifth round: Standard Eosin (Sigma-Aldrich, HT110132-1L, Lot SLCF0068) manual staining**, Standard tissue dehydration was done with gradient Ethanol (Sigma, R8382-1GA) and xylene

(Epredia, 6601). Slides were mounted with EcoMount (Biocare, EM897L). Full slide imaging was acquired at 20X (200X final magnification) with Zeiss AXIO Scanner.
